# Highly fucosylated *N*-glycans at the synaptic vesicle and neuronal plasma membrane

**DOI:** 10.1101/2022.07.06.499060

**Authors:** Mazdak M. Bradberry, Trenton M. Peters-Clarke, Evgenia Shishkova, Edwin R. Chapman, Joshua J. Coon

## Abstract

At neuronal synapses, synaptic vesicles (SVs) require glycoproteins for normal trafficking, and *N*-linked glycosylation is required for delivery of the major SV glycoproteins synaptophysin and SV2A to SVs. The molecular compositions of SV *N*-glycans, which may drive important neurobiological processes, are largely unknown. In this study, we combined organelle isolation techniques, fluorescence detection of *N*-glycans, and high-resolution mass spectrometry to characterize *N*-glycosylation at synapses and SVs from mouse brain. Detecting over 2,500 unique glycopeptides from over 550 glycoproteins, we found that abundant SV proteins harbor *N*-glycans with fucose on their complex antennae, and we identify a highly fucosylated *N*-glycan enriched in SVs as compared to synaptosomes. Antennary fucosylation was also characteristic of plasma membrane proteins and cell adhesion molecules with established roles in synaptic function and development. Our results represent the first defined *N*-glycoproteome of a neuronal organelle and raise new questions in the glycobiology of synaptic pruning and neuroinflammation.

## INTRODUCTION

Synaptic vesicles (SVs) are small recycling organelles that store and release neurotransmitters at nerve terminals. Post-translational modifications of SV proteins may play important roles in SV formation and function (Brager et al., 2003; de Jong et al., 2016; Stewart et al., 2020; Zhang et al., 2015). Among these modifications, glycosylation of luminal asparagine residues (*N*-linked glycosylation) is integral to membrane protein folding and trafficking (Cherepanova et al., 2016; Reily et al., 2019), and several of the most abundant SV proteins with established roles in neurotransmission are *N*-glycoproteins. These include the Ca^2+^ sensor synaptotagmin-1 (Bradberry et al., 2020; Geppert et al., 1994; Matthew et al., 1981), the endocytosis-related tetraspanin synaptophysin (Jahn et al., 1985; Kwon and Chapman, 2011), and the SV2A/B/C family of transporter-like glycoproteins (Bradberry and Chapman, 2022; Buckley and Kelly, 1985; Crowder et al., 1999; Custer et al., 2006). SV2A and synaptophysin both require intact glycosylation sites for normal trafficking (Kwon and Chapman, 2012), and glycosylation of SV2C is essential for the uptake of botulinum neurotoxin A (Yao et al., 2016).

Congenital disorders of glycosylation, which may disrupt the synthesis and/or degradation of *N*-linked glycans, often profoundly impact brain function (Freeze et al., 2015). As compared to other tissues, the brain contains a distinct profile of *N*-glycans in which high-mannose, oligomannose, and bisected N-acetyl-glucosamine (GlcNAc)-rich structures predominate (Ji et al., 2015; Lee et al., 2020; Shimizu et al., 1993; Williams et al., 2022). Certain glycans, including an oligosaccharide known as the Lewis^X^ antigen/SSEA-1/CD15, represent specific cell surface markers during neuronal development (Capela and Temple, 2002; Götz et al., 1996), and the brain *N*-glycome is dynamic over the lifespan (Klarić et al., 2021). Loss of the enzyme fucosyltransferase-9 (Fut9) (Kaneko et al., 1999), which generates Lewis^X^ in mouse brain by addition of antennary fucose in α(1-3) linkages to GlcNAc on complex *N*-glycans in the Golgi apparatus (Kudo et al., 2007; Nishihara, 2003), is associated with impaired neurite outgrowth *in vitro* (Gouveia et al., 2012) and behavioral abnormalities in the mouse (Kudo et al., 2007). Strikingly, the corresponding gene *FUT9* has been linked to schizophrenia in human genome-wide association studies, along with several other glycosylation-related genes (Mealer et al., 2020; Schizophrenia Working Group of the Psychiatric Genomics Consortium, 2014). Aberrant glycosylation has long been detected in the brains of patients with schizophrenia, though largely through the use of indirect methods such as immunoblotting and immunostaining (Williams et al., 2020). These findings suggest that glycobiological processes may contribute to the development of schizophrenia, but specific pathophysiologic mechanisms remain poorly understood in this context.

Recent advances in mass spectrometry-based glycoproteomics methods have enabled the detection of hundreds to thousands of unique glycopeptides obtained from mouse brain (Khidekel et al., 2007; Liu et al., 2015; Polasky et al., 2020; Riley et al., 2019; Trinidad et al., 2013). While these studies have improved our understanding of the heterogeneous distribution of *N*-glycosylation in the brain (Riley et al., 2019), the broad scope of these studies has limited their insights into roles for specific glycans in specific biological processes. Examining protein glycosylation in particular cell types or organelles, by contrast, offers an opportunity to link glycosylation with function. In particular, the abundance of glycoproteins in SVs (Bradberry et al., 2022; Takamori et al., 2006) makes these organelles attractive for the study of glycans in membrane protein trafficking. Moreover, nerve terminals represent a brain tissue component vulnerable to injury and degeneration, especially in the context of neuroinflammation (Moraes et al., 2015; Sheppard et al., 2019; Weinhard et al., 2018), schizophrenia (Onwordi et al., 2020) and age-related cognitive decline (Hark et al., 2021; Hsia et al., 1999; Terry et al., 1991). We also note a large body of evidence suggesting specialized extracellular matrix glycans at nerve terminals (Dani and Broadie, 2012). Changes in glycoprotein trafficking at the synapse may thus influence a broad range of clinically important processes. A meaningful understanding of glycan-dependent processes at the synapse, however, has been limited by the lack of a deeply characterized synaptic *N*-glycoproteome.

In this study, we establish a molecular foundation for synaptic glycobiology by defining the *N*-linked glycoproteome of SVs and their milieu. Using organelle immunoisolation techniques (Bradberry et al., 2022) and complementary analytical methods, we demonstrate that SV glycoproteins are enriched for specific glycans bearing antennary fucosylation. Highly fucosylated glycans were found primarily on SV proteins, cell adhesion molecules with known synaptic functions, and other proteins with key roles at the plasma membrane including transporters and neurotransmitter receptors. Representing the first defined glycoproteome of a secretory organelle, our results evince a common glycosylation pathway shared by plasma membrane and SV proteins and provide protein-level evidence to inform hypotheses linking protein glycosylation and synaptic biology.

## RESULTS

### Proteomic characterization of SVs and their milieu

Proteomic, glycoproteomic, and *N*-linked glycomic studies were conducted using synaptic material from whole mouse brain prepared by two approaches. Synaptosomes, which represent a “classical” crude preparation containing mostly pre- and post-synaptic elements, were prepared by differential centrifugation (**Fig. 1A**) (Takamori et al., 2006). Alongside synaptosomes, a highly pure population of SVs was isolated by immunoprecipitation (IP) with magnetic beads conjugated in-house to a monoclonal antibody against SV2 (Buckley and Kelly, 1985), according to recently described procedures (**Fig. 1A**) (Bradberry et al., 2022). These preparations were subjected to proteomic analysis by trypsin digestion and nano-flow liquid chromatography-tandem mass spectrometry (nLC-MS/MS) using an Orbitrap Eclipse mass spectrometer (Bradberry et al., 2022) (**Fig. 1B**). The synaptosome preparation was rich in cytoskeletal and mitochondrial proteins (**Fig. 1C, Supplementary Table S1**), consistent with a low degree of specific enrichment for any particular cellular compartment. In contrast, SV2-IP yielded SVs of exceptionally high purity, as evinced by the nearly exclusive presence of well-established SV proteins among the top 25 most intense in this preparation (**Fig. 1D, Table S1**). More than 2,300 proteins were detected in this SV preparation, among which over 1700 were also identified in the synaptosome fraction (**Fig. 1E**). This represents the largest number of proteins detected in a highly pure SV preparation to date (Bradberry et al., 2022; Taoufiq et al., 2020). Label-free quantitation (LFQ) intensity scores of synaptosome and SV proteins were positively correlated (**Fig. 1F**), consistent with an expected contribution from SVs to the synaptosomal proteome. Type-A gamma-amino butyric acid (GABA_A_), N-methyl-D-aspartate (NMDA), and α-amino-3-hydroxy-5-methyl-4-isoxazolepropionic acid (AMPA) receptor subunits were present in synaptosomes but undetectable in SV samples (**Fig 1G**). SV samples likewise did not contain the postsynaptic scaffolding proteins PSD-95 or gephyrin, both of which were present in synaptosome samples (**Fig. 1G**). These results establish that our synaptosome preparation contains not just mature SVs but also a more general sample of brain proteins that represents the synaptic milieu.

**Fig 1.**
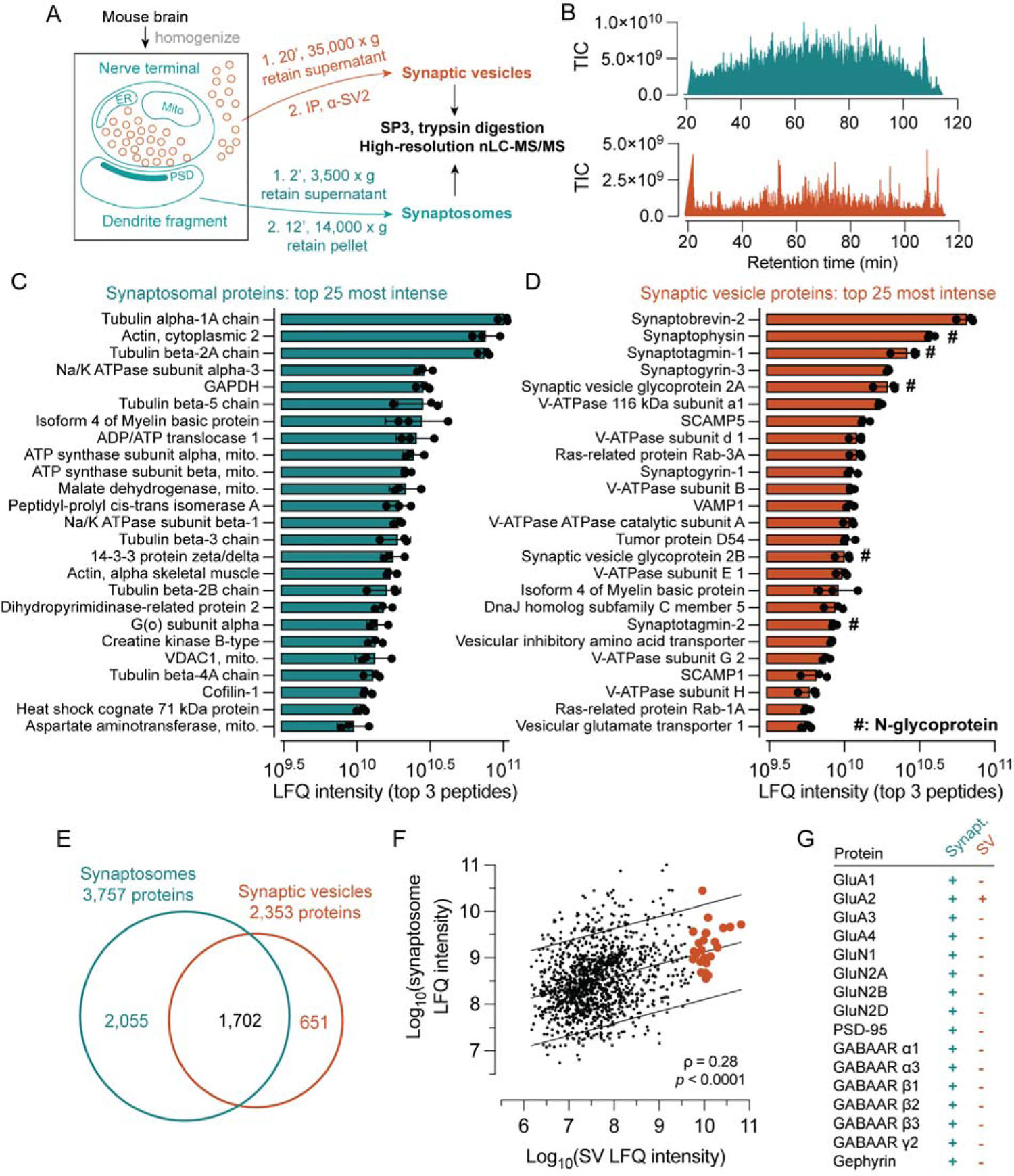
Proteomic characterization of synaptic vesicles and synaptosomes. (A) Purification scheme. SVs were isolated using magnetic beads conjugated in-house to a monoclonal α-SV2 mAb, while synaptosomes were isolated by differential centrifugation. Both sample types underwent similar processing methods and identical LC-MS/MS analysis procedures. (B) Example MS^1^ total ion current (TIC) chromatograms for SV and synaptosome samples. (C) The top 25 most intense proteins by label-free quantification analysis in synaptosome samples. Cytoskeletal and mitochondrial proteins predominated, consistent with contributions from multiple subcellular components. Error bars indicate standard deviation from three biological replicates. (D) In contrast, the top 25 most intense proteins in the SV preparation are largely well-established to reside on synaptic vesicles, with only one contaminant (myelin basic protein) observed among them. (E) Venn diagram of proteins detected in synaptosome and SV samples, demonstrating the expected high degree of overlap. (F) LFQ intensities between synaptosomal and SV proteins are positively correlated (Spearman’s rho = 0.28, *p* < 0.0001), consistent with a contribution of SVs to the synaptosomal proteome. Orange points indicate proteins listed in panel D. (G) Ionotropic glutamate and GABA receptor subunits and postsynaptic scaffolding proteins were detected in synaptosomes but largely absent from SV samples, demonstrating the purity of SVs and the inclusion of postsynaptic material in the synaptosome sample. See also Supplementary Table S1, which contains all protein-level identification and quantification data used to generate this figure.

### Glycoproteomics at the synapse

With the proteomic contents of synaptosomes and SVs established, we turned our attention to the *N*-glycosylation of proteins in these two preparations. Several studies have established that brain N-glycans comprise mostly mannosylated glycans and fucosylated complex glycans bearing bisecting GlcNAc (Chen et al., 1998; Ji et al., 2015; Lee et al., 2020; Shimizu et al., 1993; Williams et al., 2022) (**Fig. 2A**). Importantly, most mannose sugars are usually removed from the glycan before attachment of GlcNAc (Williams et al., 2022), and fucose is typically first added in an α (1,6) linkage to the *N*-linked core GlcNAc before the addition of antennary fucose in α (1,3) or α (1,4) linkages (**Fig. 2A**) (Williams et al., 2022).

**Fig. 2.**
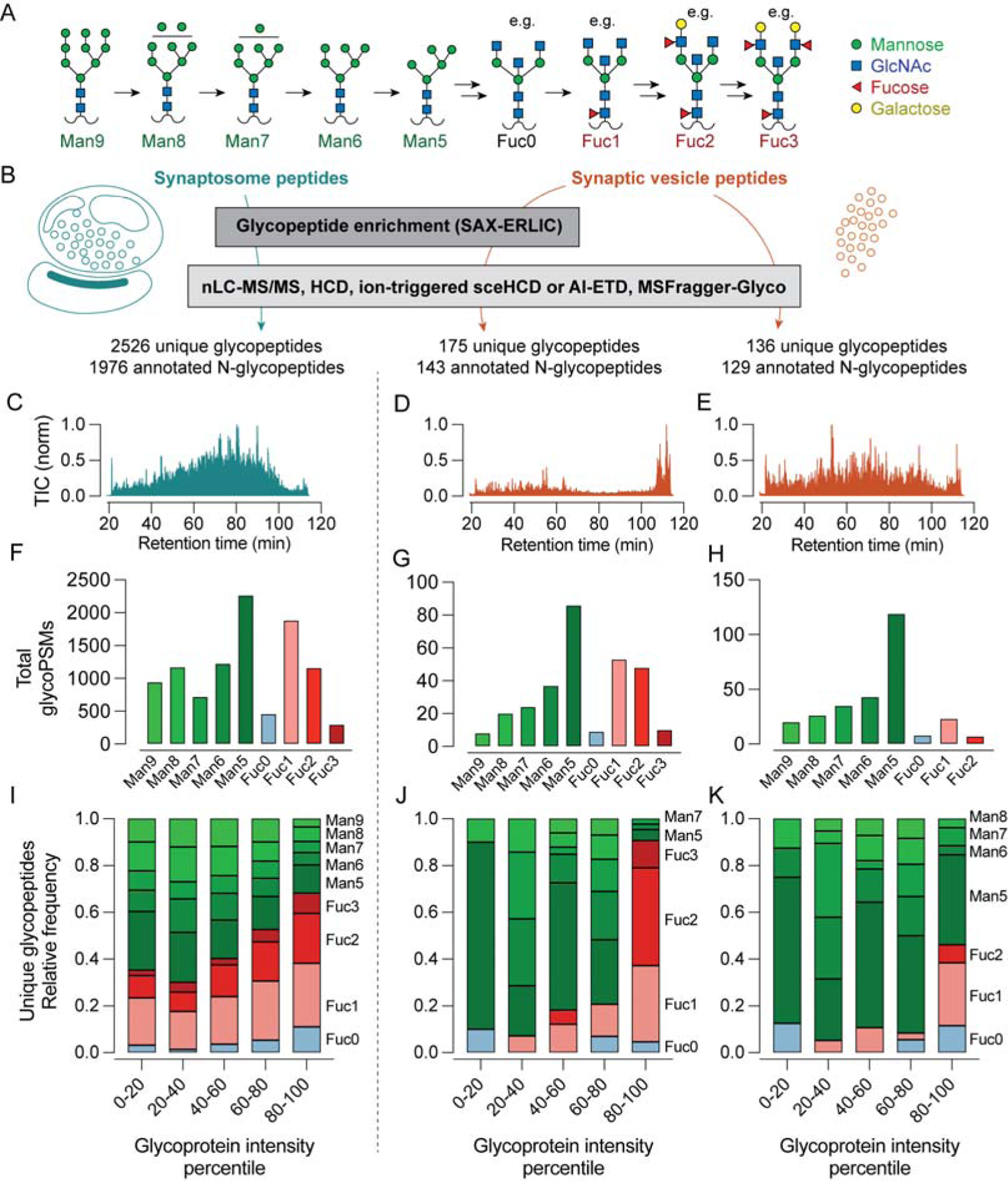
Glycoproteomics at the synapse. (A) Illustration of the *N*-glycan maturation sequence for the predominant brain *N*-glycans. In this study, glycans were annotated by their degree of mannosylation or fucosylation. Many potential glycans contain 0-3 fucoses, with major examples shown here. (B) Analysis scheme. Tryptic peptides, with or without glycan enrichment by SAX-ERLIC, were analyzed using a fragmentation strategy employing sceHCD or AI-ETD. (C-E) Example TIC chromatograms from LC-MS runs from each sample type shown above. (F-H) Distribution of all annotated glycan peptide spectral matches (glycoPSMs) identified according to the scheme in panel A. Antibody-derived glycopeptides were omitted from analysis. No Fuc3 peptides were identified in the non-enriched SV samples. (I-K) Distribution of unique annotated glycopeptides according to intensity quintile of the corresponding glycoproteins determined by standard proteomics methods (see Fig. 1). SV samples demonstrate a marked increase in Fuc2 and Fuc3 sugars in the highest quintiles, while the distribution of glycopeptides in synaptosome samples was biased to a lesser degree. See also Supplementary Table S2, containing all GlycoPSMs used to generate this figure; and Supplementary Table S3, which was used to annotate GlycoPSMs according to Man or Fuc content.

We reasoned that synaptosome and SV purification, combined with modern glycoproteomic techniques, might yield new insights into the relationship between *N*-glycosylation and SV proteins. We thus performed nLC-MS/MS experiments employing glycopeptide enrichment and Orbitrap tandem mass spectrometry with either ion-dependent stepped higher-energy collisional dissociation (sceHCD) (Riley et al., 2020) or activated-ion electron transfer dissociation (AI-ETD) (Peters-Clarke et al., 2020; Riley et al., 2019). Synaptosome samples were characterized by both AI-ETD and sceHCD-based methods after glycopeptide enrichment using strong anion exchange-electrostatic repulsion chromatography (SAX-ERLIC) prior to nLC-MS. SV samples were characterized using sceHCD both with and without glycopeptide enrichment (**Fig. 2B-E**) given the lower complexity and abundance of these samples. Importantly, SAX-ERLIC is less specific for particular *N*-glycans than lectin affinity enrichment methods (Totten et al., 2017), reducing the potential for bias towards certain glyocopeptides among enriched samples. In combination with the MSFragger/FragPipe analysis pipeline (Kong et al., 2017; Polasky et al., 2020), these data enabled a rich glycoproteomic survey of both synaptosomes and SVs purified from whole mouse brain (**Fig. 2B-E**). Glycan peptide spectral matches (GlycoPSMs) were obtained by analysis with MSFragger in glyco mode using the FragPipe analysis suite (Kong et al., 2017; Polasky et al., 2022, 2020) and filtered by excluding those glycoPSMs with a calculated false discovery rate cut-off (Q-value) of < 0.025. A total of over 2,500 unique glycopeptides from over 550 glycoproteins was identified by this method (**Fig. 2B, Supplementary Table S2**), with the majority observed in the substantially more complex synaptosome samples (**Fig. 2B**). The identified glycopeptides were annotated according to their degree of mannosylation or fucosylation (**Fig. 2A-K**) with guidance from recently published work describing the composition of the brain *N*- and *O-*glycomes (Trinidad et al., 2013; Williams et al., 2022) (**Supplementary Table S3**). GlycoPSMs containing sialic acid or unusually large numbers of sugars were omitted from annotation, as we inferred that these glycoPSMs could correspond to peptides bearing both *N*-and *O*-glycosylation on the same tryptic peptide given the low prevalence of sialylated *N*-linked glycans, and high prevalence of sialylated *O*-linked glycans, in brain (Williams et al., 2022). Despite these limitations, approximately 80% of glycoPSMs were annotated. Distributions of glycoPSMs, corresponding to major *N*-glycan types showin in **Fig. 2A,** are shown in **Fig. 2F-H**. In agreement with the work of Williams et al. (2022), we generally observed a predominance of oligomannose species along with fucosylated complex species (**Fig. 2F-H**), with a minor contribution from glycans bearing two or more fucoses. Proteins bearing three fucoses (Fuc3) were not observed in SV samples without glycopeptide enrichment (**Fig. 2H**), likely due in part to poorer positive-mode ionization efficiency of peptides bearing larger glycans.

While informative, the distribution of glycoPSMs may not reflect the actual composition of *N*-glycans in SVs or synaptosomes, as these data do not account for differences in abundance among glycoproteins giving rise to these glycopeptides. We thus analyzed these data further by considering the intensity scores obtained in our standard proteomics experiments (**Fig. 1**) for each identified glycoprotein (**Fig. 2I-K**). Glycoproteins were grouped into quintiles based on their intensity values (**Fig. 1**), and the distribution of glycoPSMs was analyzed for each quintile in each sample type (**Fig. 2I-K**). This analysis revealed a striking bias toward fucosylation in the most abundant contributors to the SV proteome (**Fig. 2J-K**), which was less pronounced in the synaptosome samples (**Fig. 2I**). This trend was particularly evident for glycans containing two or three fucoses (i.e., highly fucosylated glycans) (**Fig. 2I-K**). In glycan-enriched SV samples, nearly all the unique glycoPSMs for the most abundant SV proteins contained one or more fucoses (**Fig. 2J**). A bias towards fucosylated glycopeptides in the most abundant SV proteins was observed whether or not glycopeptide enrichment was used (**Fig. 2J-K**), demonstrating that this observation is not an artifact of the enrichment procedure. However, a stark increase in high fucosylation in abundant SV proteins was best observed using glycopeptide enrichment (**Fig. 2J**) Together, these results suggest that abundant SV proteins are rich in highly fucosylated *N*-glycans.

### Characterization of SV and synaptosome glycans by HILIC-HPLC

While the above results provide evidence that abundant SV glycoproteins are biased toward fucosylation, additional caveats exist to the interpretation of bottom-up glycoproteomics data. For example, unexpected biases in the positive-mode ionization efficiency of certain glycopeptides could cause skewed results (Harvey, 1993). We thus employed an orthogonal, ionization-independent *N*-glycan analysis method involving fluorescent detection of enzymatically cleaved glycans from SV and synaptosome samples (**Fig. 3A**). *N*-glycans were specifically and quantitatively removed from proteins with PNGase F (**Supplementary Figure S1**), followed by labeling with the fluorescent molecule procainamide via reductive amination (**Fig. 3A**). Labeled *N*-glycans were analyzed by amide hydrophilic interaction chromatography with fluorescence detection (HILIC HPLC, **Fig. 3**). Digestion with exoglycosidases specific for mannose, antennary fucose, or galactose (**Fig. 3B**) enabled a semi-quantitative determination of the contributions of various glycans to the SV and synaptosomal *N*-glycomes (**Fig. 3C-L**). The identities of the mannosylated glycans were determined using a man5 standard and glucose homopolymer ladder, defining a clear sequence of glycans bearing 5-9 mannose residues (**Supplementary Fig. S2**). Among mannosylated glycans, we observed a trend towards predominance of Man5 in both synaptosomes and SVs (**Fig. 3I**), but we did not observe significant differences between the two sample types either in the distribution of mannosylated glycans (**Fig. 3I**) or the total contribution from mannosylated glycans (**Fig. 3J**).

**Fig. 3.**
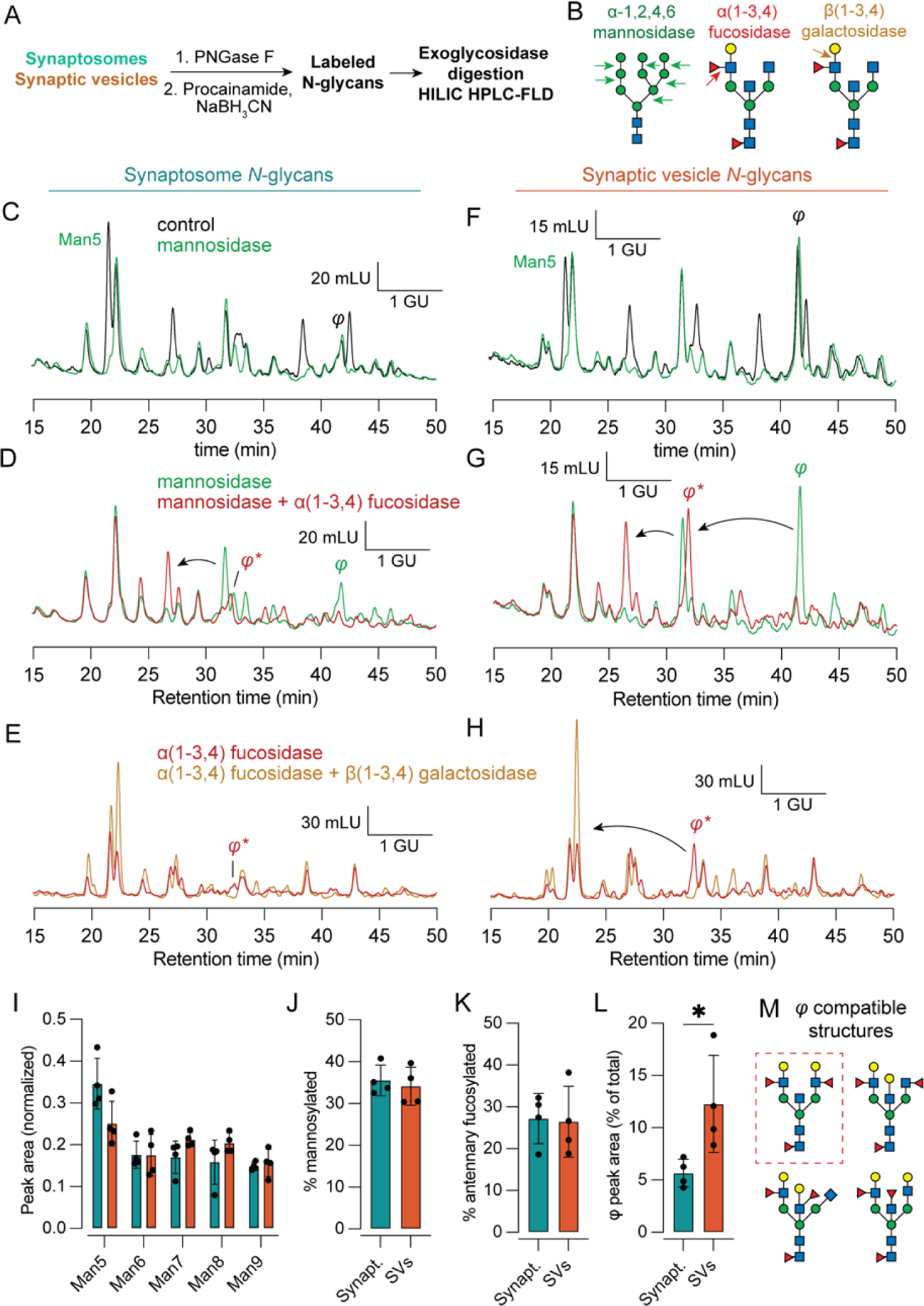
Fluorescence HPLC of released *N*-glycans defines a highly fucosylated species enriched on SVs. (A) Experimental scheme. *N*-linked glycans from SVs and synaptosomes were quantitatively released from denatured proteins using PNGase F, labeled by reductive amination with procainamide, and detected by fluorescence after separation by amide HILIC HPLC. (B) Illustration of cleavage sites for exoglycosidases used in this figure. α(1-2,4,6) mannosidase cleaves mannose residues from mannosylated *N*-glycans, α(1-3,4) fucosidase removes only antennary α (1-3,4) linked fucoses but not the core fucose, and β(1-3,4) galactosidase removes antennary galactose residues. (C) Representative chromatograms of procainamide-labeled synaptosome *N*-glycans under control conditions and treated with mannosidase. The identity of Man 5 was established using an authentic standard. Mannosidase-sensitive peaks appear in green. GU, glucose units, determined via procainamide-labeled glucose homopolymer standard; mLU, milli-luminance units. (D) The addition of α (1-3,4) fucosidase reveals the contribution from antennary fucosylated glycans in synaptosomes. (E) The combination of α (1-3,4) fucosidase and β(1-3,4) galactosidase demonstrates a modest contribution from galactosylated glycans in synaptosomes. (F, G, H) As in (C, D, E) but for SV *N*-glycans. A late-eluting α(1-3,4) fucosidase sensitive peak, denoted φ, represents a fucosylated *N*-glycan more pronounced in SVs versus synaptosomes. This peak migrates by approximately ∼1.9 GU upon α (1-3,4) fucosidase treatment, indicating the presence of two antennary fucoses. φ was further sensitive to β (1-3,4) galactosidase, which caused an additional ∼2 GU shift (panel H). φ was also subject to further cleavage by α (1-2,4,6) fucosidase, demonstrating core fucosylation (Supplementary Figure 3) (I-K) The distribution of peak areas among mannosylated glycans, as well as the total contribution from mannosylated and antennary fucosylated glycans, was equivalent in synaptosomes and SVs. (L) The peak corresponding to highly fucosylated glycan φ made a significantly larger contribution to SV versus synaptosome glycans (*p* = 0.02, Welch’s test), demonstrating enrichment of this highly fucosylated glycan in SVs. (M) Possible structures for glycan φ compatible with HPLC, glycoproteomics, and published brain *N*-glycome studies (Williams et al., 2022). Dashed box indicates a species previously predicted by Williams et al. (2022). See also Supplementary Figures S2-S4, which contain additional raw HPLC data used to establish identity of Man5-9 and φ.

In accordance with our LC-MS glycoproteomics data (**Fig. 2**), both SV and synaptosomal samples contained a major contribution from antennary fucosylated glycans (**Fig. 3D, G, K**), equal to ∼25% of all *N*-linked glycans. Strikingly, a late-eluting antennary fucosylated species bearing two antennary fucose residues, denoted φ, was significantly enriched in the SV samples (**Fig. 3D,G; Fig. 3L**). This peak was susceptible to further cleavage when incubated with α(1-2,4,6) fucosidase (**Supplementary Fig. S3**), demonstrating that this species is also core fucosylated and thus contains 3 total fucoses. This peak was also susceptible to cleavage with β(1-3,4) galactosidase, which caused another ∼2 glucose unit shift in the elution time of this peak (**Fig. 3H**). The resulting peak was sensitive to cleavage with βGlcNAcase (**Fig. S3**). Examination of our glycoproteomics data (**Fig. 2, Table S2**) demonstrated a predominant Fuc3 species, HexNAc_5_Hex_5_Fuc_3_, on the abundant SV glycoproteins SV2B and synaptophysin in glycopeptide-enriched SV samples. Several potential structures for φ, informed by our mass spectrometry data, HPLC data with enzyme digestion, and previous characterizations of the mouse brain *N*-glycome (Helm et al., 2021; Williams et al., 2022), are given in **Fig. 3M**. These structures represent complex *N*-glycans bearing up to two Lewis^X^ moieties on their antennae. We did not observe sensitivity to α(1-2) fucosidase among synaptosome or SV glycans (**Supplementary Fig. S4**), consistent with the absence of the corresponding fucosyltransferase enzymes Fut1 and Fut2 in mouse brain (Williams et al., 2022). Finally, we ensured that our findings were not due to elution of SV2 mAb from the magnetic beads, as these SV2 mAb glycans eluted at different times and were not sensitive to α(1-3,4) fucosidase orα(1-2,4,6) mannosidase (**Fig. S4**). These results confirm that both SVs and synaptosomes are rich in antennary-fucosylated glycans, with specific enrichment of at least one highly fucosylated species in SVs.

### Deep characterization of the synaptic vesicle glycoproteome

Previous work has shown that *N*-glycosylation can be remarkably heterogeneous not only across glycosites for a given protein, but also at any given glycosite (Riley et al., 2019). Moreover, while the above results support the enrichment of at least one glycan with antennary fucosylation on SV proteins, SVs still contain a substantial proportion of oligomannose glycans (**Fig. 2, 3**). A deeper examination of the glycosylation sites on each major SV glycoprotein would yield additional insights into the nature and distribution of protein glycosylation in SVs. We thus immunoprecipitated SV2 and syt1 from detergent-solubilized synaptosomes and analyzed these samples by glycan-targeted LC-MS to characterize *N*-glycosylation more thoroughly for these major SV glycoproteins (**Table S2**). The resulting data were combined with our SV glycoproteomics data (**Fig. 2, Table S2**) to define the set of unique glycopeptides present at each glycosite in each major SV glycoprotein (**Fig. 4**). In accordance with the work of Riley et al. (2019), we found that each glycosite could carry any of several unique glycans, and heterogeneity across glycosites was observed for each protein (**Fig. 4**). Strikingly, while high fucosylation was observed on each protein except for syt1, high fucosylation was usually observed on only one glycosylation site per protein (**Fig. 4**). In the case of synaptophysin, which harbors a single glycosylation site, only fucosylated glycans were detected. In SV proteins with multiple glycosylation sites, especially SV2B, non-fucosylated glycosylation sites harbored oligomannose or high-mannose glycans (**Fig. 4**). We did not observe high molecular weight forms of SV2 previously characterized by immunoblotting as keratan sulfate proteoglycans (Scranton et al., 1993) (**Supplementary Fig. S5**). Rather, this earlier result may have represented an experimental artifact, because we found that boiling SV or synaptosome samples caused an apparent increase in the molecular weight of SV2 (**Fig. S5**). Thy1, which is among the most abundant neuronal plasma membrane glycoproteins and is also found in SVs (Bradberry et al., 2022; Jeng et al., 1998; Morris, 2018), contained fucosylation at all three glycosites but was likewise biased toward fucosylation at a single site, N94 (**Fig. 4**). By contrast, syt1 was unique in its lack of fucosylation (**Fig. 4**). Indeed, the single *N*-glycosylation site on syt1 was observed to contain at least six mannose residues, even though Man5 was the most common mannosylated glycan observed in each sample (**Fig. 2, Fig. 3**) and in the brain generally (Williams et al., 2022). Syt1 is also unique among these proteins in lacking a predicted globular luminal domain. Together, these results paint a highly resolved picture of the SV glycoproteome, raise new questions about the trafficking itineraries of SV proteins, and suggest important biochemical constraints on protein glycosylation in Golgi apparatus.

**Fig. 4.**
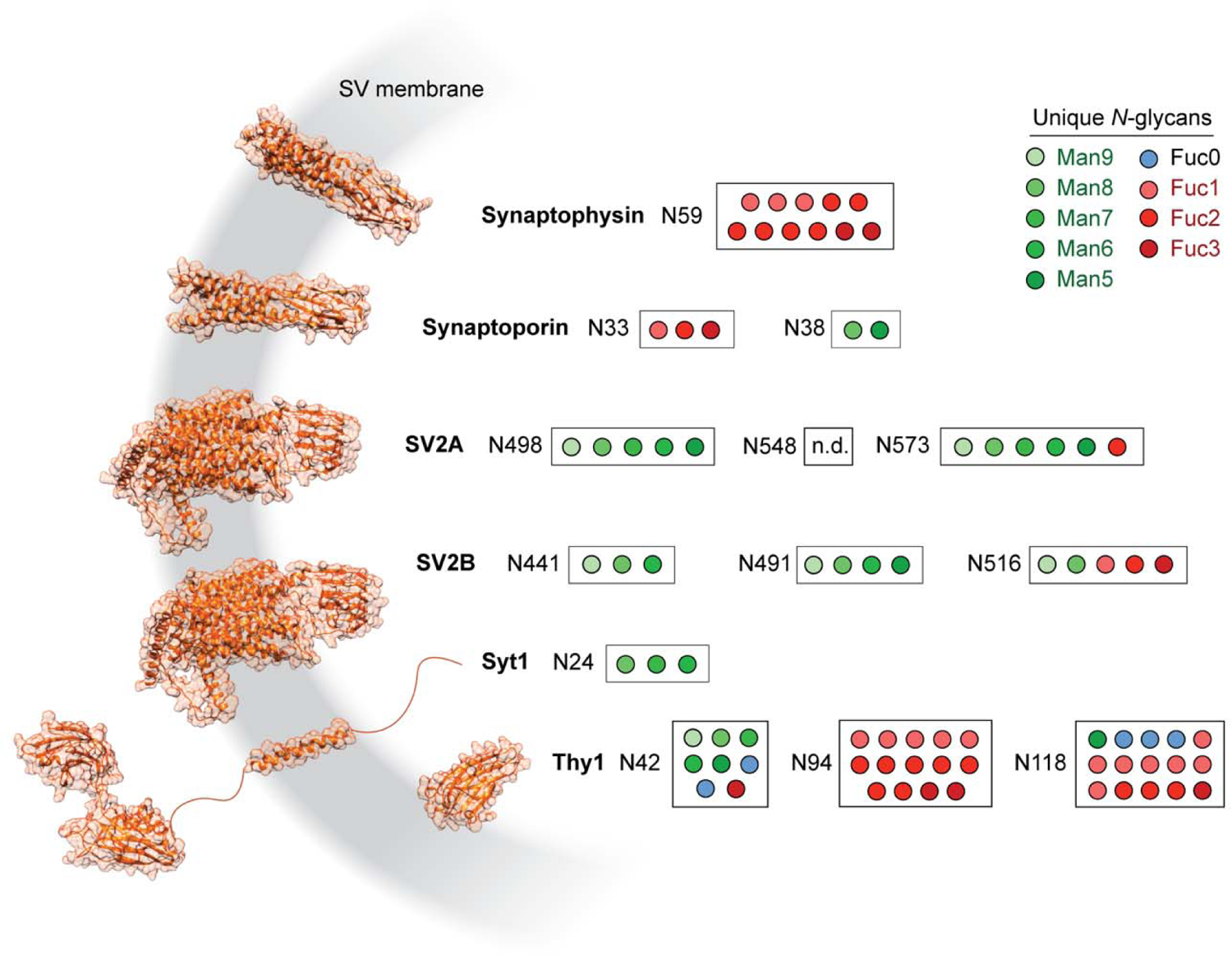
Deep glycoproteomics of the synaptic vesicle reveals site-specific biases towards mannosylation or fucosylation of SV proteins. Several of the most abundant SV glycoproteins are depicted with AlphaFold structures (Jumper et al., 2021) and symbols corresponding to unique *N*-glycopeptides detected for each *N*-linked glycosylation site. Sites are noted by their amino acid sequence number for each glycosylated asparagine. High fucosylation was observed for all SV glycoproteins shown here except for syt1. SV glycoproteins with multiple glycosylation sites demonstrated a bias towards fucosylation at a single site, while other sites were more likely to retain less mature mannosylated *N*-glycans. No glycoPSMs were observed for peptides containing N548 on SV2A. See also Supplementary Table S2, which contains the glycoPSM data used to generate this figure; and Supplementary Figure S5, which addresses prior studies on the glycosylation of SV2.

### Fucosylated *N*-glycans are characteristic of synaptic vesicle and plasma membrane proteins

We conducted further analyses of our glycoproteomics data to better define the biological context for antennary fucosylation at central synapses. Among unique glycopeptides found in SVs, the majority (164/235) were also found in synaptosome samples (**Fig. 5A**), consistent with the substantial observed overlap between the contents of these samples (**Fig. 1E-F**). A Venn diagram describing the distribution of all proteins with detected *N*-glycans is shown in **Fig. 5B**. As expected, the majority of the glycoproteins with annotated glycoPSMs in this study were mannosylated (**Fig. 2**), with a relatively smaller proportion of proteins observed with singly fucosylated and highly fucosylated complex *N*-glycans (**Fig. 5B**). In agreement with previous large-scale glycoproteomic studies of mouse brain (Riley et al., 2019), we observed that many proteins contained both mannosylated and fucosylated *N*-glycans (**Fig. 5B**).

**Fig. 5.**
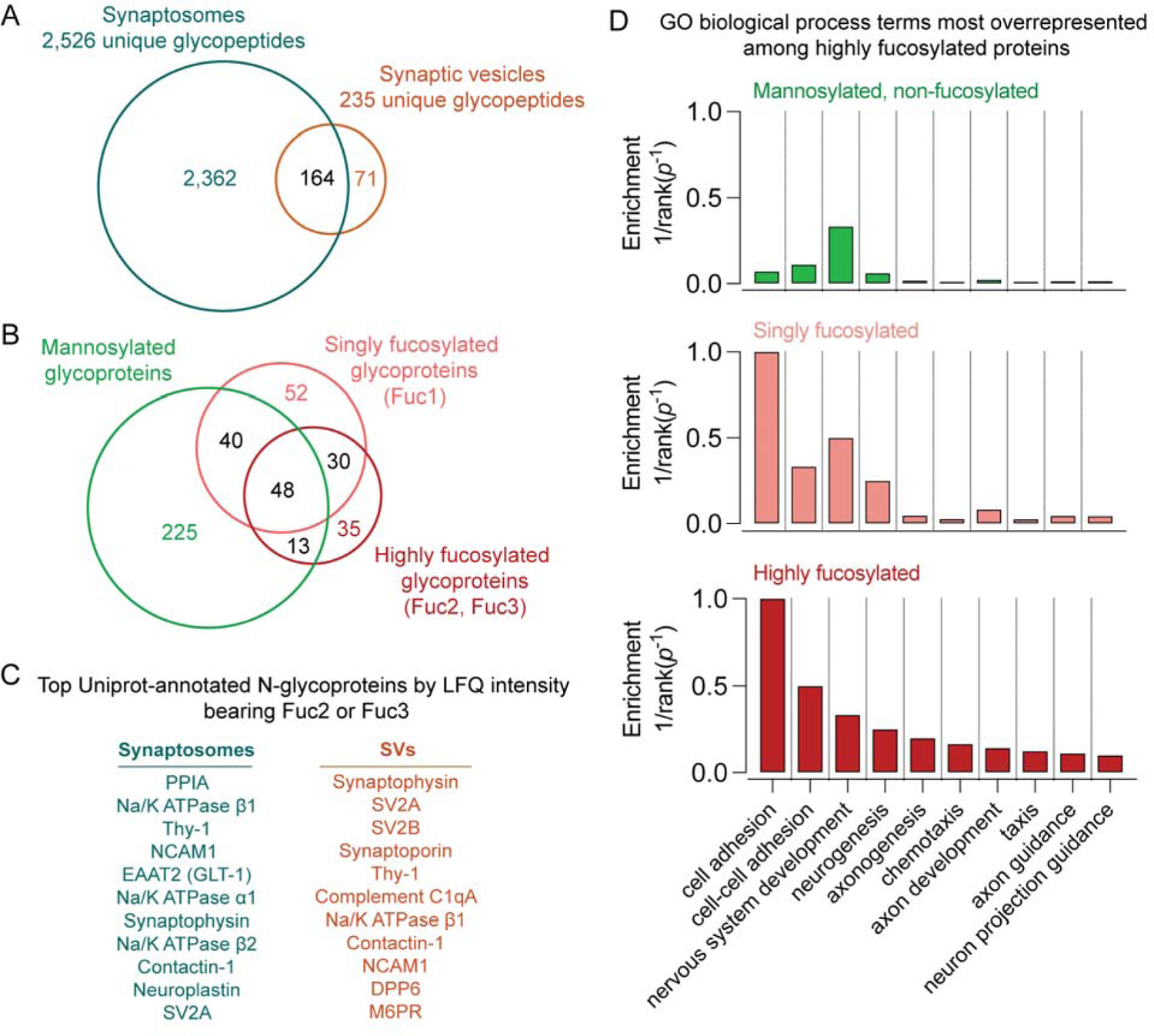
High fucosylation is characteristic of proteins at the synaptic vesicle and plasma membrane. (A) Venn diagram of unique glycopeptides detected in synaptosomes and SVs. Approximately 70% of SV glycopeptides were also detected in synaptosome samples. (B) Venn diagram of glycoproteins bearing *N*-glycans detected in this study. While some proteins were found with only one subtype of *N*-glycosylation, many proteins contained both mannosylated and fucosylated *N*-glycans. (C) The top 11 proteins by LFQ intensity with Uniprot-annotated *N*-glycosylation sites in synaptosomes and SVs that were found to bear antennary fucose, as defined by the presence of at least 2 fucoses. Antennary fucosylation was observed on almost all of the most abundant SV proteins along with a number of cell adhesion proteins with roles in synaptic development. (D) Enrichment for gene ontology (GO) biological process terms in non-fucosylated, singly fucosylated, and highly fucosylated proteins. GO terms were ranked by inverse *p*-value for enrichment, and the top 10 terms enriched in highly fucosylated proteins are shown. While substantial agreement is observed between singly fucosylated and highly fucosylated proteins, mannosylated proteins demonstrate substantially less enrichment of cell adhesion processes. See also Supplementary Table S4, containing all glycoprotein glycosylation assignments and GO biological process enrichment data used to generate this figure.

A closer examination of the identified glycopeptides in SVs and synaptosomes revealed a striking relationship among fucosylation, SV trafficking, cell adhesion, and other functions at the plasma membrane. While synaptosomes contained a substantially broader range of detected glycopeptides (**Fig. 5A**) as compared to SVs, the SV proteins synaptophysin and SV2A were among the most abundant (by LFQ intensity) highly fucosylated proteins in synaptosomes (**Fig. 5C**). Indeed, high fucosylation was observed on nearly all the most abundant SV glycoproteins, except for synaptotagmin-1/2 (**Fig. 5C, Fig. 1D, Fig. 4**). High fucosylation was also observed on cell adhesion molecules with roles in axonal pathfinding and synapse formation, including contactin-1 (Berglund et al., 1999), neuroplastin (Saito et al., 2007; Schmidt et al., 2017; the IMAGEN Consortium et al., 2015), and NCAM1 (Cremer et al., 1994) (**Fig. 5C**). The highly abundant plasma membrane glycoprotein Thy-1 (Jeng et al., 1998; Morris, 2018), the plasma membrane Na^+^/K^+^-ATPase, and the plasma membrane glutamate transporter EAAT2/GLT-1 (Furness et al., 2008; Sharma et al., 2019; Tanaka et al., 1997) were also among the top 10 most abundant highly fucosylated proteins in synaptosomes (**Fig. 5C**).

Gene ontology (GO) analysis demonstrated that cell adhesion was among the most enriched biological process terms in highly fucosylated proteins (**Fig. 5D**). Other highly enriched processes included several that are essential for synaptic development, including neurogenesis, axonogenesis, and chemotaxis (**Fig. 5D**). There was substantial overlap in top-ranked GO enrichment terms between singly fucosylated and highly fucosylated proteins (**Fig. 5D**), while proteins that were not fucosylated demonstrated substantially lower enrichment of cell adhesion-related processes (**Fig. 5D**). All ranked GO terms are shown in **Supplementary Table S4.** While a previous study using lectin affinity enrichment implied a connection between fucose-α(1,2)-galactose and a similar set of protein functions in mouse olfactory bulb (Murrey et al., 2009), we are unsure how to interpret those results given evidence that the mouse brain is largely devoid of fucose-α(1,2)-galactose and the enzymes that catalyze its addition (Murrey et al., 2009; Williams et al., 2022) (**Fig. S4**). Thus, while roles for fucosylated glycans in neurodevelopment and cell migration have previously been proposed (Gouveia et al., 2012; Kudo et al., 2007; Murrey et al., 2009), our work provides foundational evidence of links among protein fucosylation, trafficking to SVs and the plasma membrane, and cell adhesion-related processes in the mammalian brain.

## DISCUSSION

The present study demonstrates the power of combining stringent organelle purification techniques and analytical chemical methods to answer—and generate—specific questions in neurobiology. Our proteomics results (**Fig. 1**) represent coverage of the SV proteome at unprecedented depth, providing a valuable resource for investigators and extending the body of evidence supporting the use of modern IP techniques for SV purification (Bradberry et al., 2022). The combination of SV IP, glycan-focused mass spectrometry, and fluorescence methods enabled the first direct connection, to our knowledge, between a specific *N*-glycan and a brain-specific organelle (**Figs. 2-4**). We note that a prior study (Riley et al., 2019) reported a larger number of glycopeptides using whole mouse brain, lectin affinity chromatography, off-line fractionation, and activated-ion electron transfer dissociation (AI-ETD). However, a comparison between the present study and the work of Riley et al. (2019) demonstrates the utility of our focused approach using SV protein purification and SAX-ERLIC, which represents a compromise of breadth for depth at the synapse. For example, MSFragger-Glyco analysis of the dataset from Riley et al. yields no glycoPSMs for synaptophysin or synaptoporin, and the glycoPSMs found for SV2 isoforms contain at most one fucose (**Supplementary Table S5**). By contrast, we found 15 unique glycoforms for synaptophysin alone, many of which were highly fucosylated (**Fig. 4**). Moreover, while previous studies have identified brain glycans with high degrees of antennary fucosylation corresponding to Lewis^X^ or possibly fucose-α(1,2)-galactose (Chen et al., 1998; Kudo et al., 2007; Murrey et al., 2009; Trinidad et al., 2013; Williams et al., 2022), this study represents the first effort to systematically define the subcellular context for proteins bearing antennary fucosylation across the mouse brain. As with other LC-MS glycoproteomic studies (Riley et al., 2019; Trinidad et al., 2013), we do not address the large, poly-anionic proteoglycans associated with the extracellular matrix and perineuronal nets (Dani and Broadie, 2012; Testa et al., 2019), which are comparatively poor analytes for positive-mode mass spectrometry. Future application of negative ionization and fragmentation approaches, such as negative electron transfer dissociation (e.g., AI-NETD) (Riley et al., 2015), may enable the characterization of glycans not detected in the present study.

Our deep profiling of the SV *N*-glycoproteome raises several questions about the processing of *N*-glycans on proteins that undergo antennary fucosylation. Strikingly, SV proteins contained a single preferred site for fucosylation, while other sites were dominated by less mature mannosylated glycoforms (**Fig. 4**). These results are consistent with our HPLC results, which demonstrate that SVs still contain a substantial proportion of mannosylated glycans despite being enriched for antennary fucosylated structures (**Fig. 3**). Cleavage of mannose residues, and subsequent addition of GlcNAc and antennary fucose, thus appears to be site-specific. The basis for this site selectivity is unclear, though differential accessibility of glycan sites to glycan processing enzymes may play a role (Lee et al., 2014). Because only a single glycosylation site tends to undergo maturation in each SV protein (**Fig. 4**), it is tempting to speculate that glycan processing may be a rate-limiting step in the progress of some SV proteins through the Golgi. Examples of such glycan-dependent sorting pathways exist, most notably involving ER quality control mechanisms (Cherepanova et al., 2016) and mannose-6-phosphate receptors that recognize this specific glycan to recruit lysosomal proteins (Ghosh et al., 2003). While an oligomannose-, GlcNAc- or fucose-dependent Golgi transport lectin would provide an explanation for the limited maturation of all but one glycosite, more work is needed to address this hypothesis. We note that this site-specific maturation bias was less pronounced on highly abundant plasma membrane proteins such as Thy1 (**Fig. 4**) and Na^+^/K^+^-ATPase 1 (**Table S2**) (Riley et al., 2019). Top-down analyses of glycosylation across multiple sites on intact proteins may clarify the nature of the heterogeneity observed in our bottom-up studies.

The absence of fucosylation on syt1 (**Fig. 4**), a highly abundant SV glycoprotein, raises yet more questions. At least two potential explanations exist for this divergence from other SV glycoproteins: the luminal sequence of syt1, which lacks a globular domain, may disfavor certain glycan processing steps; or syt1 may not spend enough time in the Golgi compartments containing the requisite enzymes and metabolites. We note that, unlike the case for synaptophysin and SV2, trafficking of syt1 does not require *N-*glycosylation (Kwon and Chapman, 2012) but does require at least one C2 domain (Courtney et al., 2019), suggesting that syt1 sorts to SVs via a distinct molecular recognition process. Further experiments combining glycosylation site mutagenesis, enzyme manipulation, microscopy, and glycoproteomics may better define the site-specific determinants of SV protein glycosylation and their role in protein trafficking.

The enrichment of antennary fucosylated glycans on SV proteins, plasma membrane transporters and cell adhesion proteins (**Figs. 2-5**) is a key finding that merits further investigation. While antennary fucosylated glycans including Lewis^X^ have long been described as neuronal surface markers (Capela and Temple, 2002; Götz et al., 1996; Nishihara, 2003), specific roles for these glycans have been more elusive. In the brain, Fut9 is responsible for antennary fucosylation (Kudo et al., 2007; Williams et al., 2022), and deletion of Fut9 may drive mouse behavioral abnormalities (Kudo et al., 2007), cellular migration deficits (Kudo et al., 2007), and impaired neurite outgrowth (Gouveia et al., 2012). The abundance of antennary fucosylation on proteins important for cell adhesion and synaptogenesis (**Fig. 5**) suggests that this glycosylation may impact the biosynthesis, trafficking, or function of some of these proteins. The commonality of antennary fucosylation among SV and plasma membrane proteins (**Fig. 5**) is consistent with the notion that SV proteins first traffic to the plasma membrane prior to sorting to SVs (Régnier-Vigouroux et al., 1991). While high fucosylation is neither necessary nor sufficient for protein trafficking to SVs (**Fig. 4, Fig. 5**), the enrichment of the Fuc3 glycan φ on SVs (**Fig. 3**) suggests that SV recycling is associated with the presentation of fucosylated glycans at high density on the presynaptic plasma membrane. At present, the endogenous binding sites in brain for antennary fucose and Lewis^X^ are to our knowledge unknown. Further studies examining specific links among antennary fucosylation, protein trafficking, neuronal circuits, and synaptic physiology are needed to clarify the functional roles of these glycoforms.

Our findings are particularly striking given the recent identification of Fut9 as a gene locus linked to schizophrenia (Schizophrenia Working Group of the Psychiatric Genomics Consortium, 2014). Abnormal glycosylation of the antennary fucose-bearing proteins EAAT2/GLT-1 (Bauer et al., 2010) and GABA_A_ receptor subunits (**Supplementary Table S4**) (Mueller et al., 2014) has been described in schizophrenia postmortem brain studies. Altered fucosyltransferase enzyme levels have also been described in similar work (Mueller et al., 2017). Given that the specific brain functions of antennary fucosylation remain undefined, the potential links between this molecular feature and schizophrenia are numerous. For example, inefficient trafficking to axonal projections would broadly inhibit neurite outgrowth and neurotransmission, while reduced efficacy of cell-cell adhesion might reduce the stability of neuronal circuits. We note that fucose and fucosylated glycans are reportedly detectable in brain by MRI spectroscopy (Mountford et al., 2015), and their study may thus yield insights into the biology of schizophrenia and other developmental processes across the human life span.

Finally, we emphasize the need for further investigation into potential links among protein glycosylation, neuroinflammation, and synaptic pruning. Several studies have demonstrated a role for complement proteins in both synaptic pruning (Schafer et al., 2012; Stevens et al., 2007) and schizophrenia (Sekar et al., 2016), and lectins can directly activate the complement cascade (Garred et al., 2016). Among the best-known complement-activating lectins is mannose-binding lectin, which is activated by mannosylated glycans typically found on microorganisms but not on circulating glycoproteins (Garred et al., 2016). Mannose-binding lectin is a well-known mediator of inflammatory brain injury after ischemia or trauma (Neglia et al., 2020; Orsini et al., 2012) Strikingly, mannosylated glycans are common on neuronal glycoproteins (Williams et al., 2022) (**Fig. 2**) including at nerve terminals and SVs (**Figs. 2-3**). Abnormal expression or trafficking of neuronal *N*-glycans might thus modulate complement activation and synaptic pruning via the lectin pathway, particularly during periods of neuroinflammatory stress (Calcia et al., 2016; Weinhard et al., 2018). Future studies may define glycan-dependent processes in neuroinflammation-related brain injury, with a view toward new approaches for its treatment and prevention.

## Materials and Methods

### Animals

C57B/6J mice of either sex between 15 and 20 days of age were used for all experiments. All work was conducted according to protocols approved by the University of Wisconsin Institutional Animal Care and Use Committee.

### Antibody and bead preparation

Anti-SV2 mAb (SV2, DSHB) and anti-syt1 mAb (mAb 48, DSHB) were purified from ascites stocks generated prior to 2010 using protein G chromatography (Protein G Sepharose Fast Flow, Cytiva) and dialyzed extensively against phosphate-buffered saline (140 mM NaCl, 10 mM sodium phosphate buffer, pH 7.4). Single-use aliquots (200 µl, 300 µg) of mAb in PBS were kept frozen at −80 °C. Dynabeads M-270 Epoxy (14302D, Thermo Fisher) were coupled to mAb according to published procedures (Bradberry et al., 2022). 10 mg of Dynabeads stored in DMF were collected with a magnetic stand and resuspended in 200 µl borate buffer (100 mM boric acid-NaOH, pH 8.5). To this suspension was added 200 µl mAb solution, followed by 200 µl 3 M ammonium sulfate in borate buffer, with mixing by pipetting up and down after each addition. This coupling mixture was incubated with rotation at 37 °C overnight. The beads were then collected with a magnetic stand, the supernatant was discarded, and the beads were washed six times by trituration in 1 ml buffer followed by collection with the magnetic stand. The wash buffers were 500 mM NaCl, 50 mM ammonium acetate pH 4.5 and 500 mM NaCl, 50 mM Tris-HCl pH 8.0, used in an alternating manner (i.e., one buffer followed by the other, for three cycles total). The beads were then resuspended in 1 ml KPBS (145 mM KCl, 10 mM potassium phosphate buffer pH 7.4), transferred to a fresh tube, and collected on a magnetic stand. The supernatant was removed and the beads were resuspended at 30 mg/ml using 300 µl KPBS. Beads stored in this manner at 4 °C remained effective for at least 2 months.

### SV preparation

All buffers, tubes, and centrifuge rotors were cooled to 0-2 °C prior to use. One or two mice were anesthetized with isoflurane, euthanized, and the brains including cerebellum and brain stem were rapidly removed. Each brain was placed in a tight-fitting Teflon-glass Dounce homogenizer with 3.8 ml ice-cold potassium homogenization buffer (125 mM KCl, 25 mM potassium phosphate buffer, 5 mM EGTA, pH 7.4) containing protease inhibitors (cOmplete mini EDTA-free, Roche, 1 tablet / 10 ml buffer) and homogenized with ten strokes using an overhead mixer rotating at 900 RPM. The homogenate was centrifuged at 35,000 x g for 20 minutes at 2 °C. During centrifugation, 5 mg α-SV2 Dynabeads per brain were washed in KPBS and resuspended in two 2-ml microcentrifuge tubes (2.5 mg beads in 100 µl in each tube). The supernatant from each brain homogenate was added to two tubes containing Dynabeads (1.9 ml/tube), and the tubes were placed inside 50-ml conical tubes packed with ice and incubated with rotation for 25 minutes in a cold room. The supernatants were then discarded and each 2.5 mg portion of beads, corresponding to SVs from ½ mouse brain, was washed 3 times by gentle trituration in 1 ml ice-cold KPBS followed by collection on a magnetic stand. The beads were then resuspended and transferred to a fresh tube using KPBS, with beads bearing SVs from the same brain combined into the same tube (5 mg/tube), and the supernatant was removed. For proteomics and glycoproteomics studies, the beads were eluted using 50 µl 2% SDS containing 25 mM Tris-HCl pH 8.0 with heating to 50 °C for 5 minutes, and the eluates were frozen at −80 °C prior to use. For HPLC studies, the beads were eluted using 45 ul 0.5% SDS with heating to 50 °C for 5 minutes, followed by a second elution with 45 µl 2% n-β-dodecylmaltoside (DDM) (Gold Biotechnologies) for 5 minutes at room temperature. These eluates were combined and 2 µl 1 M triethylammonium bicarbonate (TEAB) was added prior to storage at −80 °C.

### Synaptosome preparation

Synaptosomes were prepared by removing and homogenizing brains as above except that the homogenization buffer contained 125 mM NaCl, 25 mM HEPES-NaOH, and 5 mM EGTA pH 7.4. The brain homogenate was centrifuged at 3,500 x g for 2 minutes, the pellet was discarded, and the supernatant transferred into two 2-ml microcentrifuge tubes and centrifuged for 12 minutes at 14,000 x g at 4 °C. The supernatant was discarded and the pellet was resuspended in 1530 µl 50 mM TEAB per 2-ml tube. 90 µl of 10% SDS and 180 µl 10% DDM (0.5% SDS, 1% DDM final) were added to each tube, which was incubated for 1 hour with rotation at 4 °C prior to aliquoting and freezing at −80 °C.

### Immunopurification of SV2 and syt1

SV2 and syt1 were immunoprecipitated from synaptosomes prepared as above but resuspended in 1.6 ml synaptosome homogenization buffer followed by the addition of 180 µl 10% DDM (∼1% DDM final) per one-half brain. The samples were incubated with rotation for 1 hour at 4 °C followed by pelleting of insoluble material by centrifugation at 20,000 x g for 20 minutes at 4 °C. The supernatants were transferred to new tubes, 3 mg α-syt1 or α-SV2 Dynabeads were added, and the tubes were incubated with rotation for 30 minutes. The beads were then washed with cold synaptosome resuspension buffer (4 x 1 ml) and eluted with 50 µl 2% SDS containing 25 mM Tris-HCl pH 8.0 with heating to 50 °C for 5 minutes. Eluates were stored frozen at −80 °C prior to use.

### Tryptic peptide preparation

For immunopurified SV, syt1, and SV2 samples, ∼60 µl bead eluate in SDS was combined with dithiothreitol (DTT, 100 mM stock) for a final concentration of 5 mM DTT and incubated at 50 °C for 25 minutes. Iodoacetamide (200 mM freshly prepared aqueous stock solution) was added to a final concentration of 15 mM and the reaction incubated in the dark at room temperature for 30 minutes. More DTT was then added (26 mM final concentration) to quench iodoacetamide. Dynabeads M-270 carboxylic acid (14305D, Thermo Fisher Scientific) were then added (2 µg/µl final concentration) and the tubes were mixed well, followed by the addition of 1 volume of absolute ethanol and brief incubation on a thermomixer (5’, 1000 RPM, 23 °C) to drive protein adsorption to the beads. The beads were washed three times with 200 µl 80% ethanol and transferred to a fresh tube with the final wash. The supernatant was removed and tryptic peptides were eluted from the beads by overnight digestion in 50 µl trypsin solution (V5111, Promega, 0.01 µg/µl in 100 mM ammonium bicarbonate) with shaking in a thermomixer (1000 RPM, 37 °C). Eluates from this step were used directly for LC-MS or subject to glycopeptide enrichment (*vide infra*). For synaptosomes, a similar procedure was followed but scaled up 10-fold, using 500 µl synaptosomal lysate as input, washes of 3 x 1 ml 80% ethanol, and elution in 250 µl trypsin solution.

### Glycopeptide enrichment

Glycopeptides were enriched by strong anion exchange-electrostatic repulsion chromatography (SAX-ERLIC) according to recently published procedures (Totten et al., 2017). Strong anion exchange columns (SOLA SAX 10 mg, Thermo Fisher) were washed with acetonitrile (3 x 1 ml), 100 mM triethylammonium acetate in water (3 x 1 ml), 1% trifluoroacetic acid (TFA) in water (3 x 1 ml), and 1% TFA in 95:5 MeCN:H_2_O. Tryptic peptides (100 µl for synaptosome samples, 50 µl for SV samples) were brought up to 95:5 MeCN:H_2_O with the addition of 19 volumes of MeCN and applied to the column twice. The column was then washed with 95:5 MeCN:H_2_O containing 1% TFA (6 x 1 ml). Glycopeptides were eluted with 50:50 MeCN:H_2_O containing 1% TFA (850 µl + 500 µl) followed by 95:5 H_2_O:MeCN containing 1% TFA (850 µl + 500 µl). The eluates were dried in a speedvac and stored at −80 °C. For each sample, 30 µl 0.2% formic acid in water was used to redissolve all dried eluates prior to LC-MS analysis.

### nLC-MS/MS

Data were collected using two systems, each comprising a hybrid Orbitrap mass spectrometer (Orbitrap Eclipse or Oribtrap Fusion Lumos, Thermo) interfaced to a nanoflow HPLC system (UltiMate 3000, Dionex) via a nanospray ionization source (Nanospray Flex, Thermo). Samples were separated using a column with integrated spray tip (PicoTip SIS, 25 cm long, 75 µm I.D.) packed in-house with C18 particles (BEH C18 1.7 µm, Waters) at ultra-high pressure (Shishkova et al., 2018) and held at 50 °C using a custom-built column heater. For standard proteomics experiments (**Fig. 1**), mobile phase A was 0.1% formic acid in H_2_O, mobile phase B was 0.1% formic acid in 80:20 MeCN:H_2_O, and peptides were separated using a two-hour gradient as follows: 0-17 min, 0-7% B; 17-102 min, 7-50% B; 102-104 min, 50-100% B; 104-108 min, 100% B; 108-110 min, 100-0% B; 110-120 min, 0% B. The flow rate was 310 nl/min, the spray voltage was 2 kV and the injection volume was 3 µl. For proteomics runs (**Fig. 1**), MS^1^ and MS^2^ scans were acquired in positive mode in the Orbitrap, and the following settings were used for MS^1^ spectra: resolution, 120,000; scan range, 400-1600 *m/z*; maximum injection time, 50 ms; AGC target, 400,000; normalized AGC target, 100%. MS^2^ spectra were acquired with the following settings: resolution, 30,000; scan range, 150-1800 *m/z*; maximum injection time, 60 ms; AGC target, 50,000; normalized AGC target, 100%; HCD collision energy, 30%. MS^1^ peaks were filtered based on the following criteria for fragmentation: charge state, 2-8; maximum intensity, 1E20, minimum intensity, 50,000. Monoisotopic precursor selection was used in peptide mode, and MS^1^ peaks were dynamically excluded for 20 seconds with a 20-ppm mass tolerance after being selected for fragmentation. For glycoproteomics runs (**Figs. 2, 4**), the same LC gradient and spray voltage were used for most experiments (see below). Glycoproteomics MS^1^ scans were obtained in positive mode every 3 seconds with the following settings: resolution, 120,000; scan range, 350-1800 *m/z*; maximum injection time, 50 ms; AGC target, 400,000; normalized AGC, target 100%. Peaks were selected for MS^2^ fragmentation from charge states 2-8 with dynamic exclusion in a ± 10 ppm window for 60 s after a single detection, with monoisotopic precursor selection enabled. For MS^2^ scans, precursors were fragmented with higher-energy collisional dissociation (HCD) with HCD energy 36%; resolution, 30,000; maximum injection time, 60 ms; AGC target, 50,000; normalized AGC target, 100%. A second fragmentation scan of the same precursor was triggered if one of the top 20 most abundant ions in the first MS^2^ spectrum was one of the following: *m/z* 204.0867, 138.0545, 366.1396, 274.0921, 292.1027, 126.055, 144.0655, 168.0654, 186.076. This second, triggered scan activated ion with either used stepped HCD (sceHCD) or activated ion-electron transfer dissociation (AI-ETD). SV samples were analyzed using sceHCD, while both sceHCD and AI-ETD were employed for synaptosome samples given their greater complexity and abundance of material for analysis. For AI-ETD, an Orbitrap Fusion Lumos mass spectrometer (Thermo Fisher Scientific, San Jose, CA) was retrofitted with a 60 W CO2 laser to allow for photoactivation (Ledvina et al., 2010; Peters-Clarke et al., 2020). For sceHCD, energies of 20%, 35%, and 50% were used (Riley et al., 2020). For AI-ETD, calibrated ETD reaction parameters were used, and the laser was operated at either 7% or 10% maximum power. All sceHCD and AI-ETD scans used the following parameters: scan range, 120-4000 *m/z*; maximum injection time, 200 ms; AGC target, 50,000; normalized AGC target, 100%. For synaptic vesicle samples, we also included glycoproteomics data from a set of runs using an alternative LC solvent system (four biological replicates total). For these runs, mobile phase A was 0.2% formic acid in H_2_O, mobile phase B was 90:10 isopropanol:acetonitrile with 0.2% formic acid and 5 mM ammonium formate, and peptides were separated using a two-hour gradient as follows: 0-13 min, 0% B; 13-18 min, 0-3% B; 18-88 min, 3-22% B; 88-100 min, 22-70% B; 100-101 min, 70-85% B; 101-105 min, 85% B; 105-106 min, 85-0% B; 106-120 min, 0% B. In this method the column was kept at 60 °C and the flow rate was 225 nl/min except during periods of high %B (88-106 min), when it was reduced to 200 nl/min.

### LC-MS/MS Data Analysis

Raw files from LC-MS runs were analyzed using the FragPipe software suite (v.17.1) with further processing of FragPipe output performed in R. All R scripts used are available via Github at https://github.com/mazbradberry/public/tree/glycoproteomics. For proteomics experiments with label-free quantification (LFQ), SV and synaptosome experiments were analyzed separately. Spectra were searched with MSFragger (v.3.4) (Kong et al., 2017) using a mouse proteome database downloaded from Uniprot on 11 October 2021 and the following settings: precursor mass tolerance, ± 20 ppm; fragment mass tolerance, ± 20 ppm, mass calibration and parameter optimization enabled; isotope error, 0/1/2; enzymatic cleavage, strict trypsin with up to 2 missed cleavages; peptide length, 7-50; peptide mass range, 500-5000 Da. Methionine oxidation and N-terminal acetylation were allowed as variable modifications and cysteine carbamidomethylation was included as a fixed modification. Validation was performed with PeptideProphet using closed search defaults for peptides and ProteinProphet for proteins. LFQ was performed with IonQuant (Yu et al., 2020) with match-between-runs, normalization, and MaxLFQ enabled. The following settings were used: feature detection m/z tolerance, ± 10 ppm; feature detection RT tolerance, 0.4 minutes; match between runs (MBR) tolerance 5 minutes, MBR ion FDR, 0.01; MBR peptide and protein FDR, 1; top 3 ions used for quantification with a minimum frequency of 0.5 and detection in at least 1 experiment. For glycoproteomics experiments, runs were grouped by sample type (SV, SV with enrichment, or synaptosomes with enrichment) prior to analysis. The same mouse protein database was used and spectra were searched with MSFragger (v.3.4) using glyco mode (Polasky et al., 2022, 2020) with the following settings: precursor mass tolerance, ± 20 ppm; fragment mass tolerance, ± 20 ppm, mass calibration and parameter optimization enabled; isotope error, 0/1/2; enzymatic cleavage, strict trypsin with up to 2 missed cleavages; peptide length, 7-50; peptide mass range, 400-5000 Da. Methionine oxidation and N-terminal acetylation were allowed as variable modifications and cysteine carbamidomethylation included as a fixed modification. The default 183 mass offsets corresponding to possible *N*-glycan compositions were included and restricted to asparagine residues. Labile modification search mode was set to nglycan and diagnostic Y ion and fragment masses were left as defaults. PTM-Shepherd was enabled, diagnostic ion search was enabled with default settings, and glycans were assigned in N-glycan mode with an FDR of 0.025, mass tolerance of ± 50 ppm, and isotope error range of −1 to 3. PeptideProphet and ProteinProphet were used for peptide and protein validation, respectively, with default settings. The same parameters were used for glycoproteomic studies of immunoprecipitated SV2 and syt1. Glycopeptide spectral matches (GlycoPSMs) were obtained from the psm output file, filtered for Q-values of ≤ 0.025, and annotated using the table shown in **Table S3**, with sialic acid-containing compositions omitted from annotation based on the likelihood that they represent combinations of *N*-and *O*-glycosylation on the same peptide (Williams et al., 2022). Tetra-fucosylated glycans and compositions tentatively identified as singly fucosylated oligomannose/high-mannose glycans were also detected but were omitted from analysis given their uncertain identification and small contribution (<5%) to the total glycan pool. Relative frequencies of unique annotated *N*-glycans and glycoprotein LFQ intensity quintiles (**Fig. 2**) were determined using R. For GO analysis (**Fig. 5**), gene lists were extracted from annotated glycoPSM tables and subjected to biological process GO term search (geneontology.org, accessed 25 April 2022) (Ashburner et al., 2000; The Gene Ontology Consortium et al., 2021).

### *N*-glycan release and labeling

For synaptosome samples, 50 µl synaptosomal lysate in 1% DDM and 0.5% SDS was combined with 50 µl 2% DDM and 5 µl 1 M DTT and heated to 50 °C for 15 minutes. For SV samples, 100 µl of SDS-DDM eluate (0.25% SDS, 1% DDM final) was combined with 5 µl 1M DTT and heated to 50 °C for 15 minutes. For each sample, 1 µl PNGase F (P0708, NEB) was then added, and the mixture was incubated at 37 °C for two hours. Each sample was allowed to cool to room temperature, combined with 50 µl of freshly prepared procainamide solution (40 mg/ml in 70:30 DMSO:acetic acid) and incubated on ice for 5 minutes. Samples were then centrifuged (20,000 x g, 10 minutes, 4 °C) and the supernatants (150 µl) transferred to fresh PCR tubes. 10 µl of sodium cyanoborohydride solution (5 M in 1 M NaOH, Sigma 296945) was added to each reaction, which was then incubated at 65 °C for two hours in a fume hood using a miniature PCR block. All steps involving sodium cyanoborohydride, including sample cleanup, were carried out in a fume hood. The samples were then combined with 20 volumes of MeCN (i.e., 75 µl sample was added to 1.5 ml MeCN) in 2-ml microcentrifuge tubes and subjected to cleanup by solid-phase extraction using a vacuum manifold. For each sample, a solid-phase extraction cartridge (OASIS HLB 30 mg, Waters) was equilibrated with 1 ml 95:5 MeCN:H_2_O, and the sample (∼3.2 ml) was applied. The cartridge was washed (2 x 1 ml 95:5 MeCN:H_2_O) and procainamide-labeled glycans eluted with 500 µl 50:50 MeCN:H_2_O. The eluates were dried using a speedvac, resuspended in Milli-Q water (200 µl for synaptosome samples, 40 µl for SV samples), and centrifuged to remove insoluble material.

### Exoglycosidase digestion and HPLC sample preparation

4 µl procainamide-labeled glycans were combined with 1 µl 10x sodium acetate – Ca^2+^ buffer (Glycobuffer 1, New England Biolabs), 1 µl 10x BSA (diluted 1:10 with Milli-Q water from a 100X stock, New England Biolabs), and 1 µl of each enzyme used as described in **Fig. 3** and supplemental figures **S2-S4**. Enzymes used included α mannosidase (P0768, New England Biolabs [NEB]), α(1-3,4) fucosidase (P0769, NEB), α(1-3,4) galactosidase (P0746, NEB), and β-GlcNAcase S (P0744, NEB). In reactions containing α-mannosidase and for all experiments shown in **Fig. 3**, 1.2 µl 10 mM zinc chloride (New England Biolabs) was included. Each reaction was brought up to 10 µl with Milli-Q water and incubated overnight at 37 °C in a PCR block. 15 µl of HPLC-grade acetonitrile was then added, the samples were centrifuged (20,000 x g, 10 minutes, 4 °C), and the supernatants stored on ice until HPLC analysis.

### HILIC-HPLC analysis

An Agilent HPLC system (Infinity 1260 Bio-inert) equipped with a fluorescence detector (Agilent 1260 FLD Spectra, 310 nm excitation, 370 nm emission), amide HILIC column (Agilent Glycan Mapping, 2.1 mm x 150 mm, 2.7 µm particle size) and manual injector was used for analysis of procainamide-labeled glycans. 10 µl glycan digest was injected to overfill a 5 µl home-cut sample loop and the column was kept at 40 °C. Mobile phase A was 100 mM ammonium formate, pH 4.4, mobile phase B was 100% acetonitrile. Samples were separated with a 90-minute gradient as follows: 0-60 min, 75-62.5% B; 60-62 min, 62.5-15% B; 62-82 min, 15% B; 82-85 min, 15-75% B; 85-90 min, 75% B. The flow rate was 0.25 ml/min except during periods of lower % B (62-85 min), when it was reduced to 0.175 ml/min. 30 minutes was allowed between runs for re-equilibration. Peak areas were determined by automatic integration in Agilent ChemStation software with the following settings: tangent skim mode, new exponential; tail peak skim height ratio, 5.00; front peak ski height ratio, 5.00; skim valley ratio, 20.00; baseline correction, advanced; peak to valley ratio, 500. Peaks eluting between 19 and 60 minutes were considered for analysis. For mannosylated peak determination, co-eluting peaks observed after mannosidase treatment were subtracted from the integrated mannosidase-sensitive peak area under control conditions.

### SDS-PAGE and immunoblot

Synaptosome and SV samples were combined with 4X SDS sample buffer containing DTT and heated to 50 °C for 15 minutes except as shown in Supplementary Fig. S5. 10 µl of prepared sample containing 6.7 µl synaptosome lysate or 1-1.5 µl SV eluate was subjected to SDS-PAGE on 4-20% gradient gels (Criterion TGX, Bio-Rad) and transferred to a PVDF membrane using a semi-dry blotting apparatus. Blots were blocked using TBS-T (150 mM NaCl, 10 mM Tris-HCl pH 7.4, 0.1% Tween-20) containing 5% nonfat dry milk and incubated overnight at 4 °C with primary antibody in TBS-T containing 1% nonfat dry milk. Antibodies for immunoblot included guinea pig anti-synaptophysin (104 211, Synaptic Systems, 1:1000 dilution of a 0.5 mg/ml stock) or mouse monoclonal anti-SV2 (SV2, DSHB, 1:1,000 dilution of a 1.2 mg/ml stock purified from ascites). Blots were washed in TBS-T, and HRP-labeled secondary antibodies were used for detection.

## Acknowledgments

We thank members of the Chapman and Coon labs for helpful discussions and feedback. This work was funded by the National Institutes of Health (grants MH061876 and NS097362 to E.R.C.; grant P41GM108538 to J.J.C.; grant T32HG002760 supporting T.M.P.C.; grant T32GM140935 supporting M.M.B). T.M.P.C. acknowledges the ACS Division of Analytical Chemistry and Agilent for support through a graduate fellowship. E.R.C. is an Investigator of the Howard Hughes Medical Institute.

## Works Cited

1. Ashburner, M., Ball, C.A., Blake, J.A., Botstein, D., Butler, H., Cherry, J.M., Davis, A.P., Dolinski, K., Dwight, S.S., Eppig, J.T., Harris, M.A., Hill, D.P., Issel-Tarver, L., Kasarskis, A., Lewis, S., Matese, J.C., Richardson, J.E., Ringwald, M., Rubin, G.M., Sherlock, G., 2000. Gene Ontology: tool for the unification of biology. Nat Genet 25, 25–29. https://doi.org/10.1038/75556

2. Bauer, D., Haroutunian, V., Meador-Woodruff, J.H., McCullumsmith, R.E., 2010. Abnormal glycosylation of EAAT1 and EAAT2 in prefrontal cortex of elderly patients with schizophrenia. Schizophrenia Research 117, 92–98. https://doi.org/10.1016/j.schres.2009.07.025

3. Berglund, E.O., Murai, K.K., Fredette, B., Sekerková, G., Marturano, B., Weber, L., Mugnaini, E., Ranscht, B., 1999. Ataxia and Abnormal Cerebellar Microorganization in Mice with Ablated Contactin Gene Expression. Neuron 24, 739–750. https://doi.org/10.1016/S0896-6273(00)81126-5

4. Bradberry, M.M., Chapman, E.R., 2022. Alleoptical monitoring of excitation–secretion coupling demonstrates that SV2A functions downstream of evoked Ca ^2+^ entry. The Journal of Physiology 600, 645–654. https://doi.org/10.1113/JP282601

5. Bradberry, M.M., Courtney, N.A., Dominguez, M.J., Lofquist, S.M., Knox, A.T., Sutton, R.B., Chapman, E.R., 2020. Molecular Basis for Synaptotagmin-1-Associated Neurodevelopmental Disorder. Neuron 107,52-64.e7. https://doi.org/10.1016/j.neuron.2020.04.003

6. Bradberry, M.M., Mishra, S., Zhang, Z., Wu, L., McKetney, J.M., Vestling, M.M., Coon, J.J., Chapman, E.R., 2022. Rapid and gentle immunopurification of brain synaptic vesicles. J. Neurosci. JN-RM-2521–21. https://doi.org/10.1523/JNEUROSCI.2521-21.2022

7. Brager, D.H., Cai, X., Thompson, S.M., 2003. Activity-dependent activation of presynaptic protein kinase C mediates post-tetanic potentiation. Nat Neurosci 6, 551–552. https://doi.org/10.1038/nn1067

8. Buckley, K., Kelly, R.B., 1985. Identification of a transmembrane glycoprotein specific for secretory vesicles of neural and endocrine cells. The Journal of Cell Biology 100, 1284–1294. https://doi.org/10.1083/jcb.100.4.1284

9. Calcia, M.A., Bonsall, D.R., Bloomfield, P.S., Selvaraj, S., Barichello, T., Howes, O.D., 2016. Stress and neuroinflammation: a systematic review of the effects of stress on microglia and the implications for mental illness. Psychopharmacology 233, 1637–1650. https://doi.org/10.1007/s00213-016-4218-9

10. Capela, A., Temple, S., 2002. LeX/ssea-1 Is Expressed by Adult Mouse CNS Stem Cells, Identifying Them as Nonependymal. Neuron 35, 865–875. https://doi.org/10.1016/S0896-6273(02)00835-8

11. Chen, Y.-J., Wing, D.R., Guile, G.R., Dwek, R.A., Harvey, D.J., Zamze, S., 1998. Neutral N-glycans in adult rat brain tissue. Complete characterisation reveals fucosylated hybrid and complex structures. Eur J Biochem 251, 691–703. https://doi.org/10.1046/j.1432-1327.1998.2510691.x

12. Cherepanova, N., Shrimal, S., Gilmore, R., 2016. N-linked glycosylation and homeostasis of the endoplasmic reticulum. Current Opinion in Cell Biology 41, 57–65. https://doi.org/10.1016/j.ceb.2016.03.021

13. Courtney, N.A., Bao, H., Briguglio, J.S., Chapman, E.R., 2019. Synaptotagmin 1 clamps synaptic vesicle fusion in mammalian neurons independent of complexin. Nat Commun 10, 4076. https://doi.org/10.1038/s41467-019-12015-w

14. Cremer, H., Lange, R., Christoph, A., Plomann, M., Vopper, G., Roes, J., Brown, R., Baldwin, S., Kraemer, P., Scheff, S., Barthels, D., Rajewsky, K., Wille, W., 1994. Inactivation of the N-CAM gene in mice results in size reduction of the olfactory bulb and deficits in spatial learning. Nature 367, 455–459. https://doi.org/10.1038/367455a0

15. Crowder, K.M., Gunther, J.M., Jones, T.A., Hale, B.D., Zhang, H.Z., Peterson, M.R., Scheller, R.H., Chavkin, C., Bajjalieh, S.M., 1999. Abnormal neurotransmission in mice lacking synaptic vesicle protein 2A (SV2A). Proceedings of the National Academy of Sciences 96, 15268–15273. https://doi.org/10.1073/pnas.96.26.15268

16. Custer, K.L., Austin, N.S., Sullivan, J.M., Bajjalieh, S.M., 2006. Synaptic Vesicle Protein 2 Enhances Release Probability at Quiescent Synapses. Journal of Neuroscience 26, 1303–1313. https://doi.org/10.1523/JNEUROSCI.2699-05.2006

17. Dani, N., Broadie, K., 2012. Glycosylated synaptomatrix regulation of trans-synaptic signaling. Devel Neurobio 72, 2–21. https://doi.org/10.1002/dneu.20891

18. de Jong, A.P.H., Meijer, M., Saarloos, I., Cornelisse, L.N., Toonen, R.F.G., Sørensen, J.B., Verhage, M., 2016. Phosphorylation of synaptotagmin-1 controls a post-priming step in PKC-dependent presynaptic plasticity. Proc Natl Acad Sci USA 113, 5095–5100. https://doi.org/10.1073/pnas.1522927113

19. Freeze, H.H., Eklund, E.A., Ng, B.G., Patterson, M.C., 2015. Neurological Aspects of Human Glycosylation Disorders. Annu. Rev. Neurosci. 38, 105–125. https://doi.org/10.1146/annurev-neuro-071714-034019

20. Furness, D.N., Dehnes, Y., Akhtar, A.Q., Rossi, D.J., Hamann, M., Grutle, N.J., Gundersen, V., Holmseth, S., Lehre, K.P., Ullensvang, K., Wojewodzic, M., Zhou, Y., Attwell, D., Danbolt, N.C., 2008. A quantitative assessment of glutamate uptake into hippocampal synaptic terminals and astrocytes: New insights into a neuronal role for excitatory amino acid transporter 2 (EAAT2). Neuroscience 157, 80–94. https://doi.org/10.1016/j.neuroscience.2008.08.043

21. Garred, P., Genster, N., Pilely, K., Bayarri-Olmos, R., Rosbjerg, A., Ma, Y.J., Skjoedt, M.-O., 2016. A journey through the lectin pathway of complement-MBL and beyond. Immunol Rev 274, 74–97. https://doi.org/10.1111/imr.12468

22. Geppert, M., Goda, Y., Hammer, R.E., Li, C., Rosahl, T.W., Stevens, C.F., Südhof, T.C., 1994. Synaptotagmin I: A major Ca2+ sensor for transmitter release at a central synapse. Cell 79, 717– 727. https://doi.org/10.1016/0092-8674(94)90556-8

23. Ghosh, P., Dahms, N.M., Kornfeld, S., 2003. Mannose 6-phosphate receptors: new twists in the tale. Nat Rev Mol Cell Biol 4, 202–213. https://doi.org/10.1038/nrm1050

24. Götz, M., Wizenmann, A., Reinhardt, S., Lumsden, A., Price, J., 1996. Selective Adhesion of Cells from Different Telencephalic Regions. Neuron 16, 551–564. https://doi.org/10.1016/S0896-6273(00)80074-4

25. Gouveia, R., Schaffer, L., Papp, S., Grammel, N., Kandzia, S., Head, S.R., Kleene, R., Schachner, M., Conradt, H.S., Costa, J., 2012. Expression of glycogenes in differentiating human NT2N neurons. Downregulation of fucosyltransferase 9 leads to decreased Lewisx levels and impaired neurite outgrowth. Biochimica et Biophysica Acta (BBA) - General Subjects 1820, 2007–2019. https://doi.org/10.1016/j.bbagen.2012.09.004

26. Hark, T.J., Rao, N.R., Castillon, C., Basta, T., Smukowski, S., Bao, H., Upadhyay, A., Bomba-Warczak, E., Nomura, T., O’Toole, E.T., Morgan, G.P., Ali, L., Saito, T., Guillermier, C., Saido, T.C., Steinhauser, M.L., Stowell, M.H.B., Chapman, E.R., Contractor, A., Savas, J.N., 2021. Pulse-Chase Proteomics of the App Knockin Mouse Models of Alzheimer’s Disease Reveals that Synaptic Dysfunction Originates in Presynaptic Terminals. Cell Systems 12, 141–158.e9. https://doi.org/10.1016/j.cels.2020.11.007

27. Harvey, D.J., 1993. Quantitative aspects of the matrix-assisted laser desorption mass spectrometry of complex oligosaccharides. Rapid Commun. Mass Spectrom. 7, 614–619. https://doi.org/10.1002/rcm.1290070712

28. Helm, J., Grünwald-Gruber, C., Thader, A., Urteil, J., Führer, J., Stenitzer, D., Maresch, D., Neumann, L., Pabst, M., Altmann, F., 2021. Bisecting Lewis X in Hybrid-Type *N*-Glycans of Human Brain Revealed by Deep Structural Glycomics. Anal. Chem. 93, 15175–15182. https://doi.org/10.1021/acs.analchem.1c03793

29. Hsia, A.Y., Masliah, E., McConlogue, L., Yu, G.-Q., Tatsuno, G., Hu, K., Kholodenko, D., Malenka, R.C., Nicoll, R.A., Mucke, L., 1999. Plaque-independent disruption of neural circuits in Alzheimer’s disease mouse models. Proceedings of the National Academy of Sciences 96, 3228– 3233. https://doi.org/10.1073/pnas.96.6.3228

30. Jahn, R., Schiebler, W., Ouimet, C., Greengard, P., 1985. A 38,000-dalton membrane protein (p38) present in synaptic vesicles. Proceedings of the National Academy of Sciences 82, 4137–4141. https://doi.org/10.1073/pnas.82.12.4137

31. Jeng, C.-J., McCarroll, S.A., Martin, T.F.J., Floor, E., Adams, J., Krantz, D., Butz, S., Edwards, R., Schweitzer, E.S., 1998. Thy-1 Is a Component Common to Multiple Populations of Synaptic Vesicles. Journal of Cell Biology 140, 685–698. https://doi.org/10.1083/jcb.140.3.685

32. Ji, I.J., Hua, S., Shin, D.H., Seo, N., Hwang, J.Y., Jang, I.-S., Kang, M.-G., Choi, J.-S., An, H.J., 2015. Spatially-Resolved Exploration of the Mouse Brain Glycome by Tissue Glyco-Capture (TGC) and Nano-LC/MS. Anal. Chem. 87, 2869–2877. https://doi.org/10.1021/ac504339t

33. Jumper, J., Evans, R., Pritzel, A., Green, T., Figurnov, M., Ronneberger, O., Tunyasuvunakool, K., Bates, R., Žídek, A., Potapenko, A., Bridgland, A., Meyer, C., Kohl, S.A.A., Ballard, A.J., Cowie, A., Romera-Paredes, B., Nikolov, S., Jain, R., Adler, J., Back, T., Petersen, S., Reiman, D., Clancy, E., Zielinski, M., Steinegger, M., Pacholska, M., Berghammer, T., Bodenstein, S., Silver, D., Vinyals, O., Senior, A.W., Kavukcuoglu, K., Kohli, P., Hassabis, D., 2021. Highly accurate protein structure prediction with AlphaFold. Nature 596, 583–589. https://doi.org/10.1038/s41586-021-03819-2

34. Kaneko, M., Kudo, T., Iwasaki, H., Ikehara, Y., Nishihara, S., Nakagawa, S., Sasaki, K., Shiina, T., Inoko, H., Saitou, N., Narimatsu, H., 1999. α,3-Fucoslytransferase IX (Fuc-TIX) is very highly conserved between human and mouse; molecular cloning, characterization and tissue distribution of human Fuc-TIX. FEBS Letters 452, 237–242. https://doi.org/10.1016/S0014-5793(99)00640-7

35. Khidekel, N., Ficarro, S.B., Clark, P.M., Bryan, M.C., Swaney, D.L., Rexach, J.E., Sun, Y.E., Coon, J.J., Peters, E.C., Hsieh-Wilson, L.C., 2007. Probing the dynamics of O-GlcNAc glycosylation in the brain using quantitative proteomics. Nat Chem Biol 3, 339–348. https://doi.org/10.1038/nchembio881

36. Klarić T.S., Salopek, M., Micek, V., Gornik Kljaić O., Lauc, G., 2021. Post-natal developmental changes in the composition of the rat neocortical N-glycome. Glycobiology 31, 636–648. https://doi.org/10.1093/glycob/cwaa108

37. Kong, A.T., Leprevost, F.V., Avtonomov, D.M., Mellacheruvu, D., Nesvizhskii, A.I., 2017. MSFragger: ultrafast and comprehensive peptide identification in mass spectrometry–based proteomics. Nat Methods 14, 513–520. https://doi.org/10.1038/nmeth.4256

38. Kudo, T., Fujii, T., Ikegami, S., Inokuchi, K., Takayama, Y., Ikehara, Y., Nishihara, S., Togayachi, A., Takahashi, S., Tachibana, K., Yuasa, S., Narimatsu, H., 2007. Mice lacking α3-fucosyltransferase IX demonstrate disappearance of Lewis x structure in brain and increased anxiety-like behaviors. Glycobiology 17, 1–9. https://doi.org/10.1093/glycob/cwl047

39. Kwon, S.E., Chapman, E.R., 2012. Glycosylation Is Dispensable for Sorting of Synaptotagmin 1 but Is Critical for Targeting of SV2 and Synaptophysin to Recycling Synaptic Vesicles. J. Biol. Chem. 287, 35658–35668. https://doi.org/10.1074/jbc.M112.398883

40. Kwon, S.E., Chapman, E.R., 2011. Synaptophysin Regulates the Kinetics of Synaptic Vesicle Endocytosis in Central Neurons. Neuron 70, 847–854. https://doi.org/10.1016/j.neuron.2011.04.001

41. Ledvina, A.R., Beauchene, N.A., McAlister, G.C., Syka, J.E.P., Schwartz, J.C., Griep-Raming, J., Westphall, M.S., Coon, J.J., 2010. Activated-Ion Electron Transfer Dissociation Improves the Ability of Electron Transfer Dissociation to Identify Peptides in a Complex Mixture. Anal. Chem. 82, 10068–10074. https://doi.org/10.1021/ac1020358

42. Lee, J., Ha, S., Kim, M., Kim, S.-W., Yun, J., Ozcan, S., Hwang, H., Ji, I.J., Yin, D., Webster, M.J., Shannon Weickert, C., Kim, J.-H., Yoo, J.S., Grimm, R., Bahn, S., Shin, H.-S., An, H.J., 2020. Spatial and temporal diversity of glycome expression in mammalian brain. Proc. Natl. Acad. Sci. U.S.A. 117, 28743–28753. https://doi.org/10.1073/pnas.2014207117

43. Lee, L.Y., Lin, C.-H., Fanayan, S., Packer, N.H., Thaysen-Andersen, M., 2014. Differential Site Accessibility Mechanistically Explains Subcellular-Specific N-Glycosylation Determinants. Front. Immunol. 5. https://doi.org/10.3389/fimmu.2014.00404

44. Liu, J., Wang, F., Mao, J., Zhang, Z., Liu, Z., Huang, G., Cheng, K., Zou, H., 2015. High-Sensitivity N-Glycoproteomic Analysis of Mouse Brain Tissue by Protein Extraction with a Mild Detergent of N-Dodecyl β-D-Maltoside. Anal. Chem. 87, 2054–2057. https://doi.org/10.1021/ac504700t

45. Matthew, W.D., Tsavaler, L.T., Reichardt, L.F., 1981. Identification of a synaptic vesicle-specific membrane protein with a wide distribution in neuronal and neurosecretory tissue. The Journal of Cell Biology 91, 257–269. https://doi.org/10.1083/jcb.91.1.257

46. Mealer, R.G., Williams, S.E., Daly, M.J., Scolnick, E.M., Cummings, R.D., Smoller, J.W., 2020. Glycobiology and schizophrenia: a biological hypothesis emerging from genomic research. Mol Psychiatry 25, 3129–3139. https://doi.org/10.1038/s41380-020-0753-1

47. Moraes, C.A., Santos, G., Spohr, T.C.L. de S. e, D’Avila, J.C., Lima, F.R.S., Benjamim, C.F., Bozza, F.A., Gomes, F.C.A., 2015. Activated Microglia-Induced Deficits in Excitatory Synapses Through IL-1β: Implications for Cognitive Impairment in Sepsis. Mol Neurobiol 52, 653–663. https://doi.org/10.1007/s12035-014-8868-5

48. Morris, R.J., 2018. Thy-1, a Pathfinder Protein for the Post-genomic Era. Front. Cell Dev. Biol. 6, 173. https://doi.org/10.3389/fcell.2018.00173

49. Mountford, C., Quadrelli, S., Lin, A., Ramadan, S., 2015. Six fucose-α(1-2) sugars and α fucose assigned in the human brain using *in vivo* two-dimensional MRS: SIX FUCOSE SUGARS ASSIGNED IN THE HUMAN BRAIN USING *IN VIVO* 2D MRS. NMR Biomed. 28, 291–296. https://doi.org/10.1002/nbm.3239

50. Mueller, T.M., Haroutunian, V., Meador-Woodruff, J.H., 2014. N-Glycosylation of GABAA Receptor Subunits is Altered in Schizophrenia. Neuropsychopharmacol 39, 528–537. https://doi.org/10.1038/npp.2013.190

51. Mueller, T.M., Yates, S.D., Haroutunian, V., Meador-Woodruff, J.H., 2017. Altered fucosyltransferase expression in the superior temporal gyrus of elderly patients with schizophrenia. Schizophrenia Research 182, 66–73. https://doi.org/10.1016/j.schres.2016.10.024

52. Murrey, H.E., Ficarro, S.B., Krishnamurthy, C., Domino, S.E., Peters, E.C., Hsieh-Wilson, L.C., 2009. Identification of the Plasticity-Relevant Fucose-α(1−2)-Galactose Proteome from the Mouse Olfactory Bulb. Biochemistry 48, 7261–7270. https://doi.org/10.1021/bi900640x

53. Neglia, L., Oggioni, M., Mercurio, D., De Simoni, M.-G., Fumagalli, S., 2020. Specific contribution of mannose-binding lectin murine isoforms to brain ischemia/reperfusion injury. Cell Mol Immunol 17, 218–226. https://doi.org/10.1038/s41423-019-0225-1

54. Nishihara, S., 2003. alpha1,3-Fucosyltransferase IX (Fut9) determines Lewis X expression in brain. Glycobiology 13, 445–455. https://doi.org/10.1093/glycob/cwg048

55. Onwordi, E.C., Halff, E.F., Whitehurst, T., Mansur, A., Cotel, M.-C., Wells, L., Creeney, H., Bonsall, D., Rogdaki, M., Shatalina, E., Reis Marques, T., Rabiner, E.A., Gunn, R.N., Natesan, S., Vernon, A.C., Howes, O.D., 2020. Synaptic density marker SV2A is reduced in schizophrenia patients and unaffected by antipsychotics in rats. Nat Commun 11, 246. https://doi.org/10.1038/s41467-019-14122-0

56. Orsini, F., Villa, P., Parrella, S., Zangari, R., Zanier, E.R., Gesuete, R., Stravalaci, M., Fumagalli, S., Ottria, R., Reina, J.J., Paladini, A., Micotti, E., Ribeiro-Viana, R., Rojo, J., Pavlov, V.I., Stahl, G.L., Bernardi, A., Gobbi, M., De Simoni, M.-G., 2012. Targeting Mannose-Binding Lectin Confers Long-Lasting Protection With a Surprisingly Wide Therapeutic Window in Cerebral Ischemia. Circulation 126, 1484–1494. https://doi.org/10.1161/CIRCULATIONAHA.112.103051

57. Peters-Clarke, T.M., Schauer, K.L., Riley, N.M., Lodge, J.M., Westphall, M.S., Coon, J.J., 2020. Optical Fiber-Enabled Photoactivation of Peptides and Proteins. Anal. Chem. 92, 12363–12370. https://doi.org/10.1021/acs.analchem.0c02087

58. Polasky, D.A., Geiszler, D.J., Yu, F., Nesvizhskii, A.I., 2022. Multiattribute Glycan Identification and FDR Control for Glycoproteomics. Molecular & Cellular Proteomics 21, 100205. https://doi.org/10.1016/j.mcpro.2022.100205

59. Polasky, D.A., Yu, F., Teo, G.C., Nesvizhskii, A.I., 2020. Fast and comprehensive N- and O-glycoproteomics analysis with MSFragger-Glyco. Nat Methods 17, 1125–1132. https://doi.org/10.1038/s41592-020-0967-9

60. Régnier-Vigouroux, A., Tooze, S.A., Huttner, W.B., 1991. Newly synthesized synaptophysin is transported to synaptic-like microvesicles via constitutive secretory vesicles and the plasma membrane. The EMBO Journal 10, 3589–3601. https://doi.org/10.1002/j.1460-2075.1991.tb04925.x

61. Reily, C., Stewart, T.J., Renfrow, M.B., Novak, J., 2019. Glycosylation in health and disease. Nat Rev Nephrol 15, 346–366. https://doi.org/10.1038/s41581-019-0129-4

62. Riley, N.M., Hebert, A.S., Westphall, M.S., Coon, J.J., 2019. Capturing site-specific heterogeneity with large-scale N-glycoproteome analysis. Nat Commun 10, 1311. https://doi.org/10.1038/s41467-019-09222-w

63. Riley, N.M., Malaker, S.A., Driessen, M.D., Bertozzi, C.R., 2020. Optimal Dissociation Methods Differ for *N* - and *O*-Glycopeptides. J. Proteome Res. 19, 3286–3301. https://doi.org/10.1021/acs.jproteome.0c00218

64. Riley, N.M., Matthew J.P., R., Rose, C.M., Richards, A.L., Kwiecien, N.W., Bailey, D.J., Hebert, A.S., Westphall, M.S., Coon, J.J., 2015. The Negative Mode Proteome with Activated Ion Negative Electron Transfer Dissociation (AI-NETD). Molecular & Cellular Proteomics 14, 2644–2660. https://doi.org/10.1074/mcp.M115.049726

65. Saito, A., Fujikura-Ouchi, Y., Kuramasu, A., Shimoda, K., Akiyama, K., Matsuoka, H., Ito, C., 2007. Association study of putative promoter polymorphisms in the neuroplastin gene and schizophrenia. Neuroscience Letters 411, 168–173. https://doi.org/10.1016/j.neulet.2006.08.042

66. Schafer, D.P., Lehrman, E.K., Kautzman, A.G., Koyama, R., Mardinly, A.R., Yamasaki, R., Ransohoff, R.M., Greenberg, M.E., Barres, B.A., Stevens, B., 2012. Microglia Sculpt Postnatal Neural Circuits in an Activity and Complement-Dependent Manner. Neuron 74, 691–705. https://doi.org/10.1016/j.neuron.2012.03.026

67. Schizophrenia Working Group of the Psychiatric Genomics Consortium, 2014. Biological insights from 108 schizophrenia-associated genetic loci. Nature 511, 421–427. https://doi.org/10.1038/nature13595

68. Schmidt, N., Kollewe, A., Constantin, C.E., Henrich, S., Ritzau-Jost, A., Bildl, W., Saalbach, A., Hallermann, S., Kulik, A., Fakler, B., Schulte, U., 2017. Neuroplastin and Basigin Are Essential Auxiliary Subunits of Plasma Membrane Ca2+-ATPases and Key Regulators of Ca2+ Clearance. Neuron 96, 827–838.e9. https://doi.org/10.1016/j.neuron.2017.09.038

69. Scranton, T.W., Iwata, M., Carlson, S.S., 1993. The SV2 Protein of Synaptic Vesicles Is a Keratan Sulfate Proteoglycan. J Neurochem 61, 29–44. https://doi.org/10.1111/j.1471-4159.1993.tb03535.x

70. Sekar, A., Bialas, A.R., de Rivera, H., Davis, A., Hammond, T.R., Kamitaki, N., Tooley, K., Presumey, J., Baum, M., Van Doren, V., Genovese, G., Rose, S.A., Handsaker, R.E., Schizophrenia Working Group of the Psychiatric Genomics Consortium, Daly, M.J., Carroll, M.C., Stevens, B., McCarroll, S.A., 2016. Schizophrenia risk from complex variation of complement component 4. Nature 530, 177–183. https://doi.org/10.1038/nature16549

71. Sharma, A., Kazim, S.F., Larson, C.S., Ramakrishnan, A., Gray, J.D., McEwen, B.S., Rosenberg, P.A., Shen, L., Pereira, A.C., 2019. Divergent roles of astrocytic versus neuronal EAAT2 deficiency on cognition and overlap with aging and Alzheimer’s molecular signatures. Proc. Natl. Acad. Sci. U.S.A. 116, 21800–21811. https://doi.org/10.1073/pnas.1903566116

72. Sheppard, O., Coleman, M.P., Durrant, C.S., 2019. Lipopolysaccharide-induced neuroinflammation induces presynaptic disruption through a direct action on brain tissue involving microglia-derived interleukin 1 beta. J Neuroinflammation 16, 106. https://doi.org/10.1186/s12974-019-1490-8

73. Shimizu, H., Ochiai, K., Ikenaka, K., Mikoshiba, K., Hase, S., 1993. Structures of N-linked sugar chains expressed mainly in mouse brain. Journal of Biochemistry 114, 334–338.

74. Shishkova, E., Hebert, A.S., Westphall, M.S., Coon, J.J., 2018. Ultra-High Pressure (>30,000 psi) Packing of Capillary Columns Enhancing Depth of Shotgun Proteomic Analyses. Anal. Chem. 90, 11503–11508. https://doi.org/10.1021/acs.analchem.8b02766

75. Stevens, B., Allen, N.J., Vazquez, L.E., Howell, G.R., Christopherson, K.S., Nouri, N., Micheva, K.D., Mehalow, A.K., Huberman, A.D., Stafford, B., Sher, A., Litke, A.M., Lambris, J.D., Smith, S.J., John, S.W.M., Barres, B.A., 2007. The Classical Complement Cascade Mediates CNS Synapse Elimination. Cell 131, 1164–1178. https://doi.org/10.1016/j.cell.2007.10.036

76. Stewart, L.T., Abiraman, K., Chatham, J.C., McMahon, L.L., 2020. Increased O-GlcNAcylation rapidly decreases GABAAR currents in hippocampus but depresses neuronal output. Sci Rep 10, 7494. https://doi.org/10.1038/s41598-020-63188-0

77. Takamori, S., Holt, M., Stenius, K., Lemke, E.A., Grønborg, M., Riedel, D., Urlaub, H., Schenck, S., Brügger, B., Ringler, P., Müller, S.A., Rammner, B., Gräter, F., Hub, J.S., De Groot, B.L., Mieskes, G., Moriyama, Y., Klingauf, J., Grubmüller, H., Heuser, J., Wieland, F., Jahn, R., 2006. Molecular Anatomy of a Trafficking Organelle. Cell 127, 831–846. https://doi.org/10.1016/j.cell.2006.10.030

78. Tanaka, K., Watase, K., Manabe, T., Yamada, K., Watanabe, M., Takahashi, K., Iwama, H., Nishikawa, T., Ichihara, N., Kikuchi, T., Okuyama, S., Kawashima, N., Hori, S., Takimoto, M., Wada, K., 1997. Epilepsy and Exacerbation of Brain Injury in Mice Lacking the Glutamate Transporter GLT-1. Science 276, 1699–1702. https://doi.org/10.1126/science.276.5319.1699

79. Taoufiq, Z., Ninov, M., Villar-Briones, A., Wang, H.-Y., Sasaki, T., Roy, M.C., Beauchain, F., Mori, Y., Yoshida, T., Takamori, S., Jahn, R., Takahashi, T., 2020. Hidden proteome of synaptic vesicles in the mammalian brain. Proc Natl Acad Sci USA 117, 33586–33596. https://doi.org/10.1073/pnas.2011870117

80. Terry, R.D., Masliah, E., Salmon, D.P., Butters, N., DeTeresa, R., Hill, R., Hansen, L.A., Katzman, R., 1991. Physical basis of cognitive alterations in alzheimer’s disease: Synapse loss is the major correlate of cognitive impairment. Ann Neurol. 30, 572–580. https://doi.org/10.1002/ana.410300410

81. Testa, D., Prochiantz, A., Di Nardo, A.A., 2019. Perineuronal nets in brain physiology and disease. Seminars in Cell & Developmental Biology 89, 125–135. https://doi.org/10.1016/j.semcdb.2018.09.011

82. The Gene Ontology Consortium, Carbon, S., Douglass, E., Good, B.M., Unni, D.R., Harris, N.L., Mungall, C.J., Basu, S., Chisholm, R.L., Dodson, R.J., Hartline, E., Fey, P., Thomas, P.D., Albou, L.-P., Ebert, D., Kesling, M.J., Mi, H., Muruganujan, A., Huang, X., Mushayahama, T., LaBonte, S.A., Siegele, D.A., Antonazzo, G., Attrill, H., Brown, N.H., Garapati, P., Marygold, S.J., Trovisco, V., dos Santos, G., Falls, K., Tabone, C., Zhou, P., Goodman, J.L., Strelets, V.B., Thurmond, J., Garmiri, P., Ishtiaq, R., Rodríguez-López, M., Acencio, M.L., Kuiper, M., Lægreid, A., Logie, C., Lovering, R.C., Kramarz, B., Saverimuttu, S.C.C., Pinheiro, S.M., Gunn, H., Su, R., Thurlow, K.E., Chibucos, M., Giglio, M., Nadendla, S., Munro, J., Jackson, R., Duesbury, M.J., Del-Toro, N., Meldal, B.H.M., Paneerselvam, K., Perfetto, L., Porras, P., Orchard, S., Shrivastava, A., Chang, H.-Y., Finn, R.D., Mitchell, A.L., Rawlings, N.D., Richardson, L., Sangrador-Vegas, A., Blake, J.A., Christie, K.R., Dolan, M.E., Drabkin, H.J., Hill, D.P., Ni, L., Sitnikov, D.M., Harris, M.A., Oliver, S.G., Rutherford, K., Wood, V., Hayles, J., Bähler, J., Bolton, E.R., De Pons, J.L., Dwinell, M.R., Hayman, G.T., Kaldunski, M.L., Kwitek, A.E., Laulederkind, S.J.F., Plasterer, C., Tutaj, M.A., Vedi, M., Wang, S.-J., D’Eustachio, P., Matthews, L., Balhoff, J.P., Aleksander, S.A., Alexander, M.J., Cherry, J.M., Engel, S.R., Gondwe, F., Karra, K., Miyasato, S.R., Nash, R.S., Simison, M., Skrzypek, M.S., Weng, S., Wong, E.D., Feuermann, M., Gaudet, P., Morgat, A., Bakker, E., Berardini, T.Z., Reiser, L., Subramaniam, S., Huala, E., Arighi, C.N., Auchincloss, A., Axelsen, K., Argoud-Puy, G., Bateman, A., Blatter, M.-C., Boutet, E., Bowler, E., Breuza, L., Bridge, A., Britto, R., Bye-A-Jee, H., Casas, C.C., Coudert, E., Denny, P., Estreicher, A., Famiglietti, M.L., Georghiou, G., Gos, A., Gruaz-Gumowski, N., Hatton-Ellis, E., Hulo, C., Ignatchenko, A., Jungo, F., Laiho, K., Le Mercier, P., Lieberherr, D., Lock, A., Lussi, Y., MacDougall, A., Magrane, M., Martin, M.J., Masson, P., Natale, D.A., Hyka-Nouspikel, N., Orchard, S., Pedruzzi, I., Pourcel, L., Poux, S., Pundir, S., Rivoire, C., Speretta, E., Sundaram, S., Tyagi, N., Warner, K., Zaru, R., Wu, C.H., Diehl, A.D., Chan, J.N., Grove, C., Lee, R.Y.N., Muller, H.-M., Raciti, D., Van Auken, K., Sternberg, P.W., Berriman, M., Paulini, M., Howe, K., Gao, S., Wright, A., Stein, L., Howe, D.G., Toro, S., Westerfield, M., Jaiswal, P., Cooper, L., Elser, J., 2021. The Gene Ontology resource: enriching a GOld mine. Nucleic Acids Research 49, D325–D334. https://doi.org/10.1093/nar/gkaa1113

83. the IMAGEN Consortium, Desrivières, S., Lourdusamy, A., Tao, C., Toro, R., Jia, T., Loth, E., Medina, L.M., Kepa, A., Fernandes, A., Ruggeri, B., Carvalho, F.M., Cocks, G., Banaschewski, T., Barker, G.J., Bokde, A.L.W., Büchel, C., Conrod, P.J., Flor, H., Heinz, A., Gallinat, J., Garavan, H., Gowland, P., Brühl, R., Lawrence, C., Mann, K., Martinot, M.L.P., Nees, F., Lathrop, M., Poline, J.-B., Rietschel, M., Thompson, P., Fauth-Bühler, M., Smolka, M.N., Pausova, Z., Paus, T., Feng, J., Schumann, G., 2015. Single nucleotide polymorphism in the neuroplastin locus associates with cortical thickness and intellectual ability in adolescents. Mol Psychiatry 20, 263– 274. https://doi.org/10.1038/mp.2013.197

84. Totten, S.M., Feasley, C.L., Bermudez, A., Pitteri, S.J., 2017. Parallel Comparison of N-Linked Glycopeptide Enrichment Techniques Reveals Extensive Glycoproteomic Analysis of Plasma Enabled by SAX-ERLIC. J. Proteome Res. 16, 1249–1260. https://doi.org/10.1021/acs.jproteome.6b00849

85. Trinidad, J.C., Schoepfer, R., Burlingame, A.L., Medzihradszky, K.F., 2013. N- and O-Glycosylation in the Murine Synaptosome. Molecular & Cellular Proteomics 12, 3474–3488. https://doi.org/10.1074/mcp.M113.030007

86. Weinhard, L., di Bartolomei, G., Bolasco, G., Machado, P., Schieber, N.L., Neniskyte, U., Exiga, M., Vadisiute, A., Raggioli, A., Schertel, A., Schwab, Y., Gross, C.T., 2018. Microglia remodel synapses by presynaptic trogocytosis and spine head filopodia induction. Nat Commun 9, 1228. https://doi.org/10.1038/s41467-018-03566-5

87. Williams, S.E., Mealer, R.G., Scolnick, E.M., Smoller, J.W., Cummings, R.D., 2020. Aberrant glycosylation in schizophrenia: a review of 25 years of post-mortem brain studies. Mol Psychiatry 25, 3198–3207. https://doi.org/10.1038/s41380-020-0761-1

88. Williams, S.E., Noel, M., Lehoux, S., Cetinbas, M., Xavier, R.J., Sadreyev, R.I., Scolnick, E.M., Smoller, J.W., Cummings, R.D., Mealer, R.G., 2022. Mammalian brain glycoproteins exhibit diminished glycan complexity compared to other tissues. Nat Commun 13, 275. https://doi.org/10.1038/s41467-021-27781-9

89. Yao, G., Zhang, S., Mahrhold, S., Lam, K., Stern, D., Bagramyan, K., Perry, K., Kalkum, M., Rummel, A., Dong, M., Jin, R., 2016. N-linked glycosylation of SV2 is required for binding and uptake of botulinum neurotoxin A. Nat Struct Mol Biol 23, 656–662. https://doi.org/10.1038/nsmb.3245

90. Yu, F., Haynes, S.E., Teo, G.C., Avtonomov, D.M., Polasky, D.A., Nesvizhskii, A.I., 2020. Fast Quantitative Analysis of timsTOF PASEF Data with MSFragger and IonQuant. Molecular & Cellular Proteomics 19, 1575–1585. https://doi.org/10.1074/mcp.TIR120.002048

91. Zhang, N., Gordon, S.L., Fritsch, M.J., Esoof, N., Campbell, D.G., Gourlay, R., Velupillai, S., Macartney, T., Peggie, M., van Aalten, D.M.F., Cousin, M.A., Alessi, D.R., 2015. Phosphorylation of Synaptic Vesicle Protein 2A at Thr84 by Casein Kinase 1 Family Kinases Controls the Specific Retrieval of Synaptotagmin-1. Journal of Neuroscience 35, 2492–2507. https://doi.org/10.1523/JNEUROSCI.4248-14.2015

