## Supplemental Figures for "Highly fucosylated *N*-glycans at the synaptic vesicle and neuronal plasma membrane"

**Supplementary Information**
**Bradberry et al., 2022**

Supplementary Tables (attached)

1. Table S1: Proteomics with label-free quantification results
2. Table S2: GlycoPSMs from all samples in this study
3. Table S3: Glycan annotation table
4. Table S4: Glycoproteins and Gene Ontology enrichment analyses
5. Table S5: GlycoPSMs from analysis of a prior large-scale brain *N*-glycome study (Riley et al., 2019)

Supplementary Figures (below)

1. Figure S1: Immunoblot confirmation of deglycosylation by PNGase F
2. Figure S2: Establishment of glucose unit calibration and mannosylated glycan identities
3. Figure S3: Characterization of the SV-enriched glycan $\varphi$
4. Figure S4. SV glycan samples contain minimal contamination from eluted antibodies, and synaptic glycans are not sensitive to α(1-2) fucosidase
5. Figure S5: The apparent molecular weight of SV2 increases with application of heat

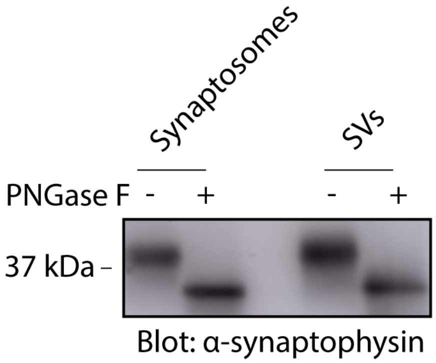


**Supplementary Figure S1. Immunoblot confirmation of deglycosylation by PNGase F**. Synaptosome and SV samples were subject to deglycosylation with PNGase F according to conditions used for glycan release and labeling (see main Fig. 3) and subjected to SDS-PAGE followed by immunoblot with anti-synaptophysin antibody. Deglycosylation of this synaptic vesicle glycoprotein causes an increase in SDS-PAGE mobility. Following treatment with PNGase F, all synaptophysin has been converted to the higher-mobility species, indicating quantitative deglycosylation for both SVs and synaptososomes under these conditions.

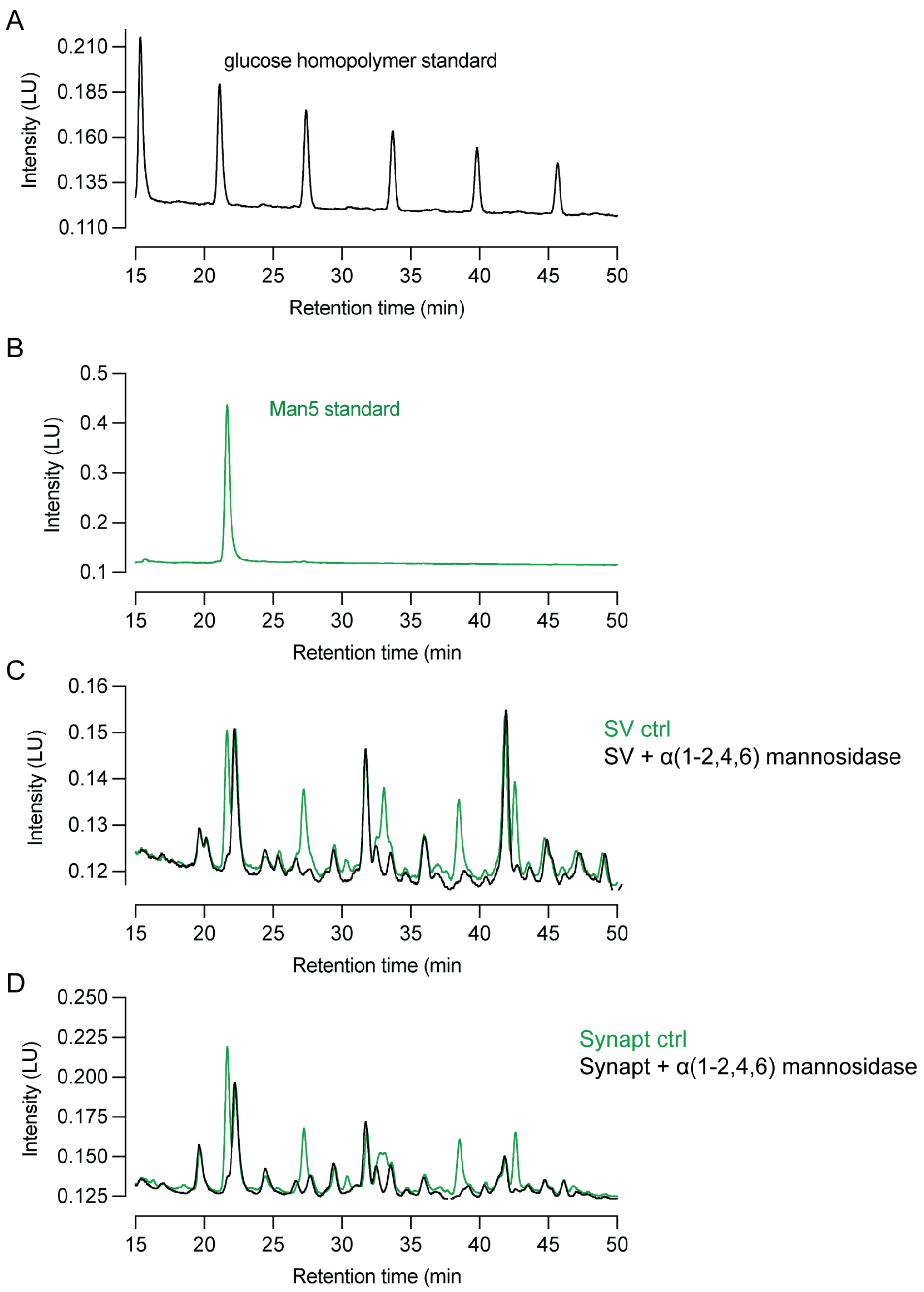


**Supplementary Figure S2. Establishment of glucose unit calibration and mannosylated glycan identities.** (A) A linear glucose homopolymer standard was labeled with procainamide and analyzed by HILIC HPLC. Individual peaks correspond to glucose polymers of increasing length, with each additional glucose in the pictured peaks causing an increase in retention time by 6.1 ± 0.2 (mean ± SD) minutes. (B) HILIC-HPLC trace of a sample containing authentic Man5 standard labeled with procainamide. (C) HILIC-HPLC trace of SV *N*-glycan sample labeled with procainamide with or without treatment with α(1-2,4,6) mannosidase. Mannosidase-sensitive peaks occur at the same position as the Man5 standard and at evenly spaced increasing retention times, consistent with well-established *N*-glycan structures containing up to 9 mannose residues in a branching configuration. (D) as in (C) but with synaptosome glycans.

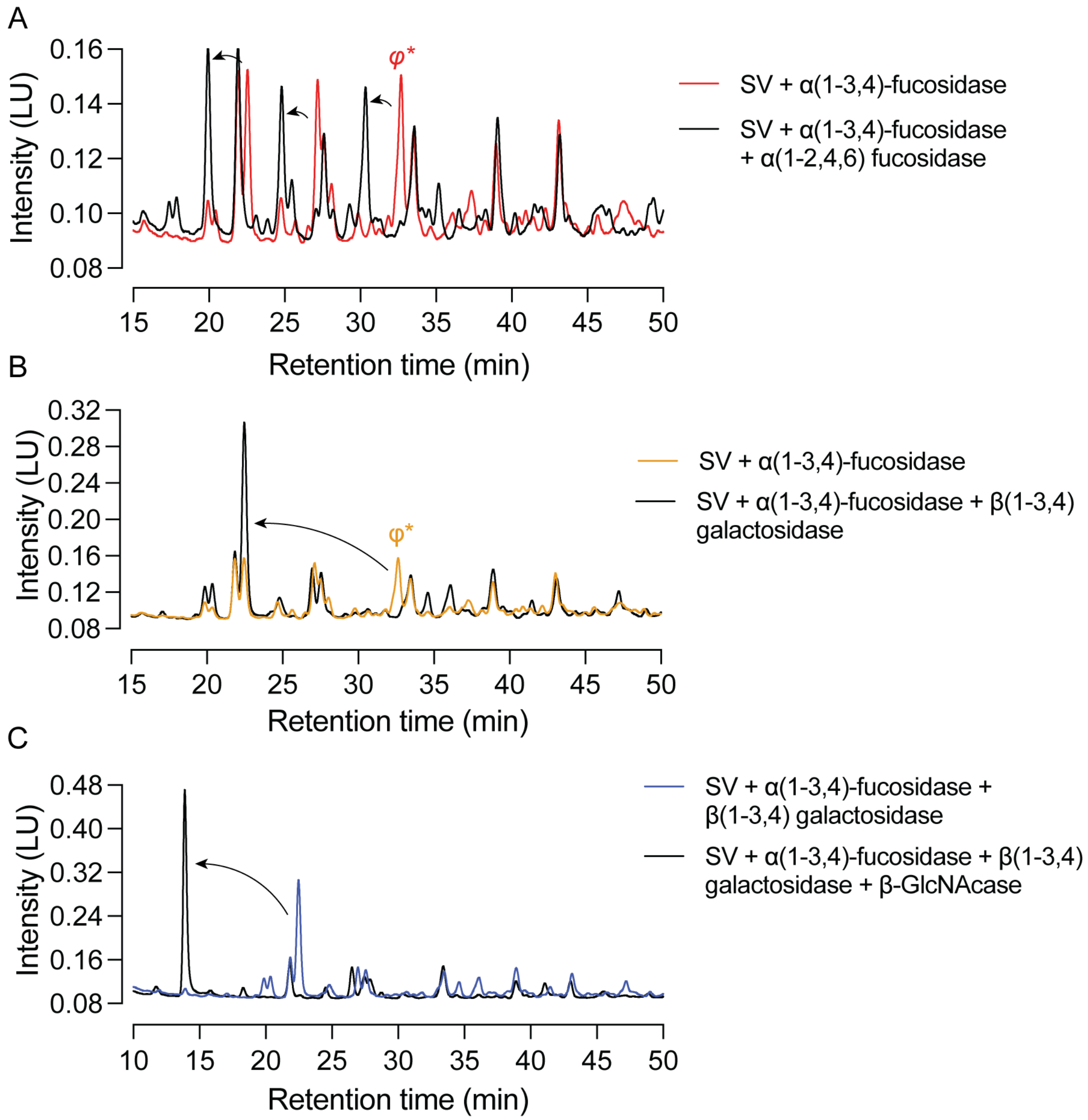


**Supplementary Figure S3. Characterization of the SV-enriched glycan *φ*** (A) φ*, the peak resulting from cleavage of antennary fucose from φ, is further sensitive to α(1-2,4,6) fucosidase cleavage, indicating the presence of core fucose not sensitive to α(1-3,4) fucosidase. (B) φ* is sensitive to β(1-3,4) galactosidase and elutes ~2 glucose units earlier after digestion with this enzyme, demonstrating the presence of two galactose sugars. (C) The peak resulting from galactosidase treatment is further sensitive to cleavage by β-GlcNAcase, confirming that φ is an elaboration of a GlcNAc-bearing complex *N*-glycan.

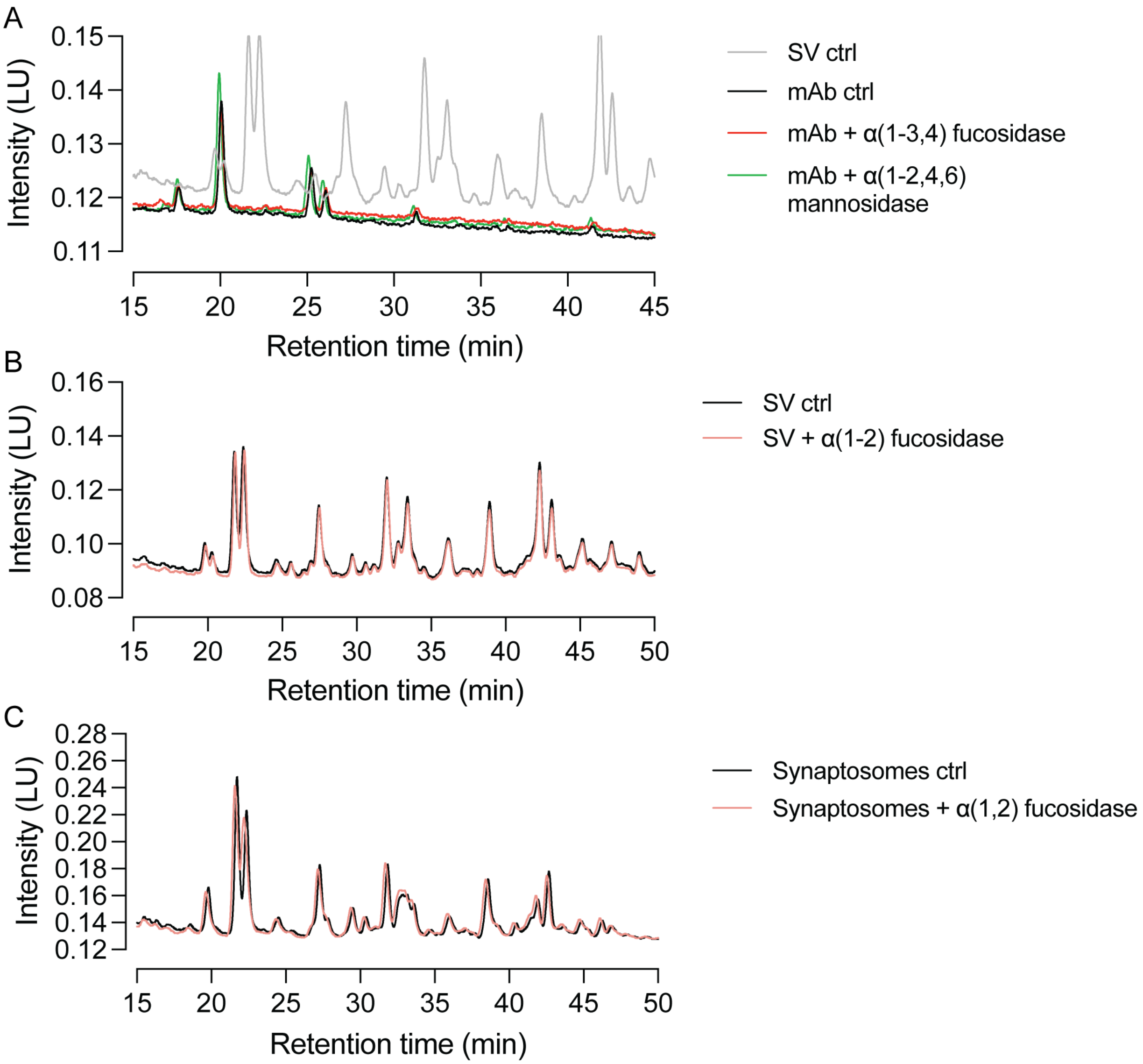


**Supplementary Figure S4. SV glycan samples contain minimal contamination from eluted antibodies, and synaptic glycans are not sensitive to α(1-2) fucosidase**. (A) HILIC-HPLC traces of procainamide-labeled SV *N*-glycans overlaid with traces from procainamide-labeled SV2 mAb *N*-glycans. SV2 mAb *N*-glycans are insensitive to α(1-3,4) fucosidase and α(1-2,4,6) mannosidase and elute at times distinct from major SV *N*-glycans. (B) α(1-2) fucosidase does not cause shifts in elution time of SV glycans, consistent with a minimal contribution of fucose α(1-2) galactose sugars to SV glycans and the absence of the enzymes (Fut1, Fut2) that catalyze the addition of fucose at the 2-position of galactose. (C) as in (B) but for synaptosome *N*-glycans.

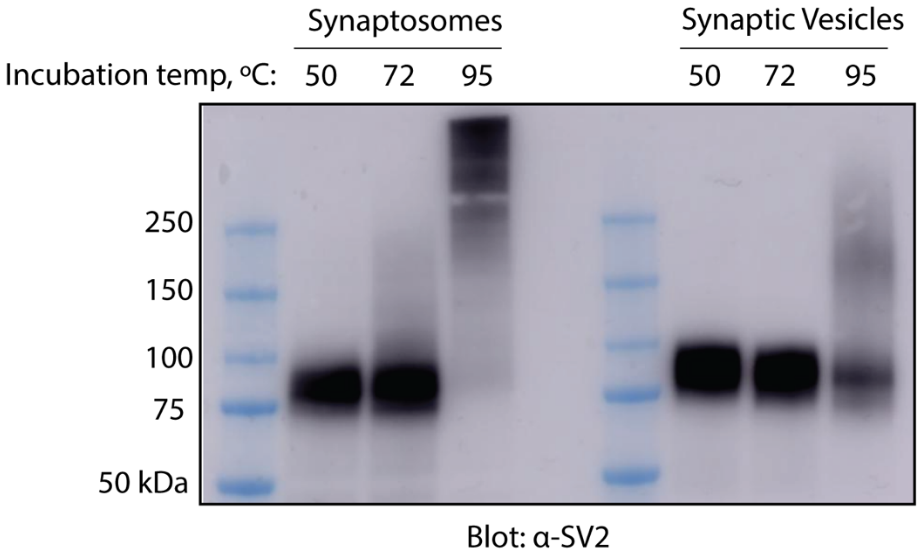


**Supplementary Figure S5: The apparent molecular weight of SV2 increases with application of heat.** Synaptosome or SV samples were combined with SDS sample buffer containing DTT and heated for 15 minutes prior to SDS-PAGE and immunoblot with anti-SV2 antibody. SV2 immunoreactivity shifted to higher molecular weight in a graded fashion with the application of increased heat.
